# Shifted PAMs generate DNA overhangs and enhance SpCas9 post-catalytic complex dissociation

**DOI:** 10.1101/2022.11.08.515552

**Authors:** Jinglong Wang, Julien Le Gall, Richard L Frock, Terence R Strick

## Abstract

*Streptococcus pyogenes* CRISPR-Cas9 (SpCas9) stabilizes each strand of a DNA bubble via distinct interactions, forming a stable ~20 bp R-loop on the complementary strand but relying on narrower protein-DNA interactions to bind the non-complementary strand with 5’ NGG protospacer adjacent motif (PAM). The enzyme’s HNH endonuclease domain cleaves DNA within the R-loop, and its RuvC endonuclease cleaves 5’ to the NGG and opposite the HNH cleavage site to generate blunt DNA ends. We show that this nucleoprotein complex can be mechanically strained by shifting the RNA:DNA hybrid and that the HNH cleavage site tracks this shift but the RuvC cleavage site does not, resulting in overhanging DNA ends. This is observed using cleavage in living cells and sequencing assays to characterize overhangs generated by stressed complexes, as well as single-molecule cleavage assays to characterize differential cleavage kinetics of HNH and RuvC endonucleases in stressed complexes as well as complex disassembly.

Using Sanger sequencing and high-throughput genome sequencing of DNA cleavage reactions, we find that the SpCas9 complex responds to internal mechanical strain by robustly generating overhanging, rather than blunt, DNA ends. Internal mechanical strain is generated by increasing or decreasing the spacing between the RNA:DNA hybrid and the downstream canonical PAM. Up to two-base 3’ overhangs can be robustly generated via a two-base increase in the distance between hybrid and PAM. We also use single-molecule experiments to reconstruct the full course of the CRISPR-SpCas9 reaction in real-time, monitoring and quantifying the R-loop formation, the first and second DNA incision events, and the post-catalytic complex dissociation. Complex dissociation and release of broken DNA ends appears to be a rate-limiting step of the reaction, and strained SpCas9 are sufficiently destabilized so as to rapidly dissociate after formation of broken DNA ends.

## Introduction

CRISPR is a bacterial and archaeal adaptive immune mechanism which mobilizes the Cas proteins and guide RNAs to interact, recognize and cleave invasive DNA (Barrangou et al., 2007; Brouns et al., 2008; Garneau et al., 2010; Marraffini et al., 2010). *Streptococcus pyogenes* (SpCas9), a well-characterized member of the type II-A Crispr-Cas family, has been widely applied toward genome editing in prokaryotes and eukaryotes (Jinek et al., 2012; Cong et al., 2013; Hsu et al., 2014). SpCas9 is a single protein of about 160 kDa with divalent ion-dependent HNH and RuvC endonuclease domains. SpCas9 can bind dual guide RNAs (dgRNAs), which are composed of the CRISPR RNA (crRNA) and the trans-activating Crispr RNA (tracrRNA) that pair to form a ribonucleoprotein (RNP). The tracrRNA is composed of 86 nts with conserved sequence and facilitates the maturation of the ~40 nt crRNA (Deltcheva et al., 2011). The 5’ region of tracrRNA and 3’ region of crRNA pair via conserved sequence to form dgRNA, and in practice, the tracrRNA and crRNA can be engineered to form single guide RNA (sgRNA) (Jinek et al., 2012). The ~20 nts spacer of the RNA guides displace the “non-complementary” DNA strand, and its hybridization with the protospacer of “complementary” DNA strand results in formation of an R-loop. The PAM interaction domain (PID) of Cas9 (Anders et al., 2014) that scans for NGG PAMs on the non-complementary strand was also found to be crucial for R-loop initiation and extension (Sternberg et al., 2014, 2015; Gong et al., 2017). During R-loop formation, the targeted DNA and dgRNA-SpCas9 undergo dramatic and ordered conformational changes (Sternberg et al., 2015; Yang et al., 2018). The HNH endonuclease domain aligns with the R-loop at the beginning of pre-catalytic complex formation between DNA and dgRNA-SpCas9, whereas the RuvC endonuclease domain becomes positioned on the non-complementary strand facing the HNH site later in the maturation of the catalytic complex (Jiang et al., 2016).

Early structural and single-molecule studies have revealed the importance of the NGG PAM in regulating SpCas9 catalytic activity (Sternberg et al., 2014), as well as poor tolerance of nucleotide mismatches between the bases of guide RNA that seed the R-loop (Hsu et al., 2013). On the other hand, the PID can be engineered to display a certain level of plasticity- NGAN, NGNG, NGCG and NAAG PAM variants were characterized in SpCas9 mutants (Hirano et al., 2016; Anders et al., 2016) and off-target DSBs formed by wildtype (WT) SpCas9 correlate to PAM variants such as GAG or AAG (Frock et al., 2015; Kleinstiver et al., 2015).

Here, we find that SpCas9 responds to offsets of several nucleotides in the spacing between the crRNA:DNA hybrid and PAM by generating overhanging DSBs, and that this “shift-PAM” targeting is enhanced by physiological levels of DNA supercoiling. We reach this conclusion by examining DNA end structures formed in ensemble biochemical and cell assays and by using single-molecule experiments to interrogate the SpCas9 catalytic mechanism: R-loop formation, first and second DNA incision events, and dissociation of the post-cleavage complex from DNA.

## Results

### Shift-PAM-targeted SpCas9 robustly cleaves linear and supercoiled DNA

We deployed canonical and shift-PAM guide RNAs to induce cleavage within a synthetic gene (Graves et al., 2015). We first designed a canonical crRNA targeting a 20 nt protospacer with a consensus NGG PAM located on the coding strand of the synthetic gene (S1_T0 crRNA). We then designed 20 nt shift-PAM crRNAs targeting DNA in base-pair increments upstream of the canonical S1_T0 crRNA target (S1_T1, S1_T2, S1_T3, and S1_T4 crRNAs). To control for cleavage we also examined two additional canonically-targeted SpCas9 sites on the non-coding strand (S2_T0 and S3_T0) (Figure 1a, Extended Data Fig.1a). The various crRNAs were hybridized with conserved tracrRNAs to form dgRNAs and subsequently combined with SpCas9 to form enzymatic dgRNA SpCas9 RNP complexes. To assess DNA strand cleavage activities in the presence or absence of supercoiling, the differentially-targeted dgRNA-SpCas9s were added in ensemble assays to either supercoiled or SbfI-linearized ~5 kb double-stranded DNA substrates and then assessed for single- or double-strand cleavage using gel electrophoresis (see Methods).

**Figure 1.**
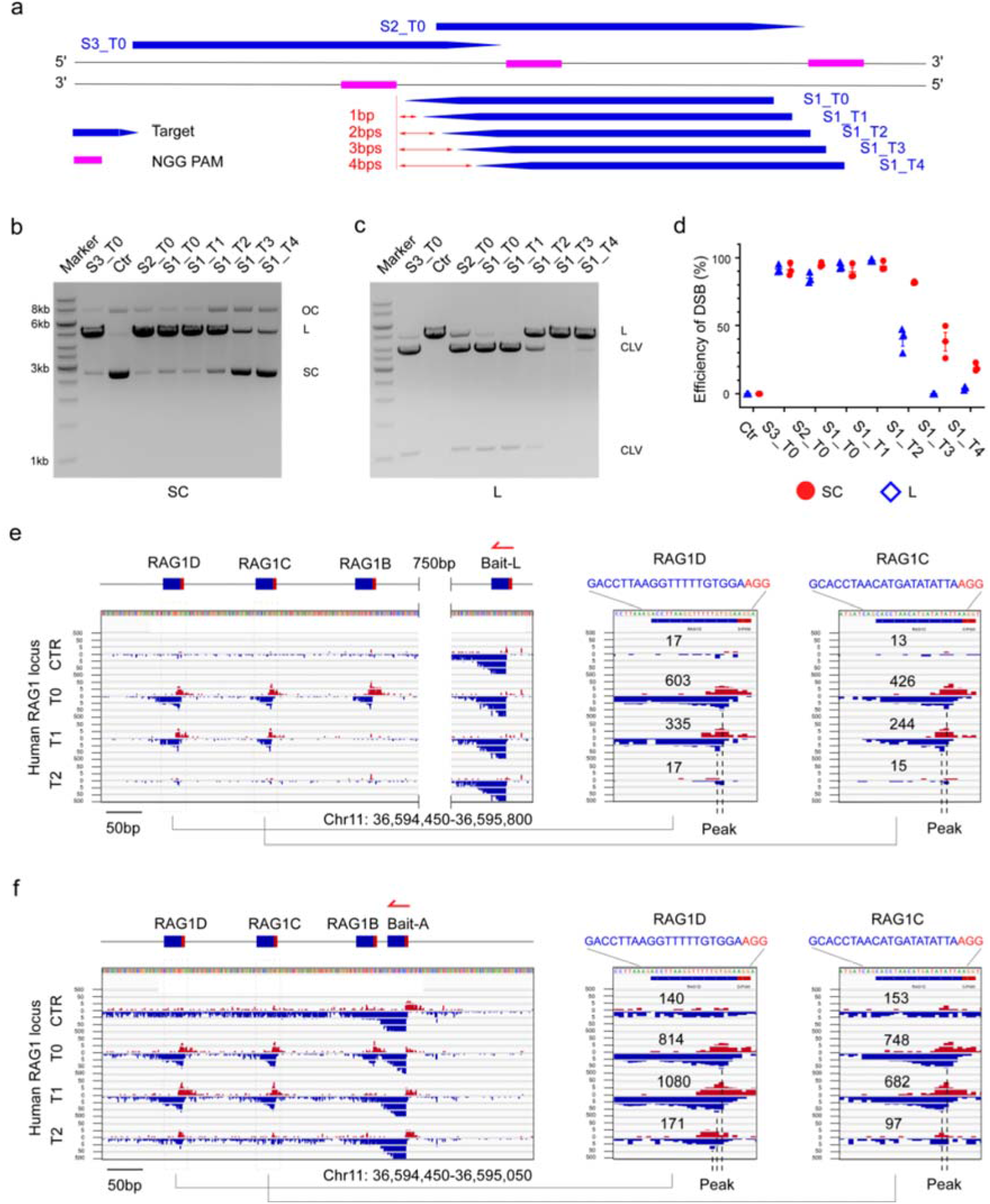
SpCas9 cleavage obtained by varying the spacing between hybridized target and PAM. **(a)** DNA loci targeted by RNPs with canonical (S1_T0 SpCas9, S2_T0 SpCas9, S3_T0 SpCas9) and shift-PAM spacing (S1_T1 SpCas9, S1_T2 SpCas9, S1_T3 SpCas9, S1_T4 SpCas9) between hybridized target and PAM. Blue arrows represent the 20 nt targeting region for each RNP and magenta rectangles represent the NGG PAM. **(b)** RNPs with canonical and shift-PAM target-PAM spacing process targeted supercoiled DNA (SC) first into nicked open circular DNA (OC) and then into linear DNA (L). **(c)** As in prior panel but with linear DNA substrate (L) processed into two cleaved products (CLV). **(d)** The normalized efficiencies of quantified cleavage of circular DNA (red dot) and linear DNA (blue diamond), with error bars (SEM). **(e-f)** LAM-HTGTS IGV plots in the RAG1 locus of HEK293T cells using two different T0-PAM guides as bait (red harpoon), Bait-L (e) and Bait-A (f), to obtain the DSB profiles in the sites RAG1B, RAG1C and RAG1D generated by CTR (bait alone), guides T0s, T1s and T2s, respectively. Enlarged figures of RAG1D and RAG1C targets (dashed boxes on left) with junction numbers are displayed in the right panels. Dashed lines indicate the peaks for the T0s, T1s, T2s targets. Plots are displayed in customized 0-20-200 scale for plus (red) and minus (blue) chromosome orientations. See Extended Table 2 for total library junctions and replication for each bait.

As predicted, canonically-targeted RNPs (S1_T0, S2_T0, and S3_T0) efficiently cleaved both DNA strands of supercoiled and linear DNA substrates within 3 hours (Figure 1b-d). Unexpectedly, shift-PAM RNPs (S1_T1 and S1_T2) generated robust levels of DSBs for supercoiled substrates, comparable to levels observed on canonically-targeted controls (Figure 1b, d). Double-strand cleavage efficiency progressively decreased with increasing shift-PAM spacing, with an increasing ratio of incised (nicked) versus linearized fragments from S1_T3 and S1_T4 RNPs (Figure 1b,d), reflecting an accumulation of partially processed intermediates. In contrast to supercoiled substrates, the decline in cleavage efficiency of the linearized substrate was much more pronounced: S1_T1 SpCas9 DSB cleavage of linear DNA was indistinguishable from that of supercoiled DNA, whereas S1_T2 SpCas9 DSB cleavage was substantially reduced and very little cleavage was detected for S1_T3 and S1_T4 RNPs (Figure 1c-d). Overall, the increased spacing between the PAM and the RNA-targeted sequences was generally more tolerated on supercoiled DNA than on linearized DNA, in a manner reminiscent of the way in which mutations in the spacer are better tolerated on negatively supercoiled DNA (Ivanov et al., 2020).

To get a better sense of how shift-PAM cleavage varies across sequence contexts, we targeted supercoiled and linear DNA at three additional sites using crRNAs implementing 2 bp spacing (S4_T2, S5_T2, S6_T2) (Extended Data Fig.1a). S5_T2 SpCas9 complexes acted robustly on supercoiled DNA substrate but much less so for linear DNA substrate, as observed with S1_T2 SpCas9. The S4_T2 and S6_T2 RNPs displayed extreme opposite activities, with the former robustly generating DSBs in both supercoiled and linear contexts and the latter only slightly nicking supercoiled DNA (Extended Data Fig.1b-d). We also considered whether SpCas9 may cleave DNA strands when the 20 nt target sequence was moved closer to the PAM in base-pair increments. Thus, we also generated S1_T-1 and S1_T-2 crRNAs (Extended Data Fig.1a) and assayed them in the ensemble assays described above. For supercoiled substrates we found robust DSB generation with S1_T-1 SpCas9 and low DSB generation with S1_T-2 SpCas9, and both of these activities were attenuated in the linear substrate ensemble (Extended Data Fig.1b-d). Collectively, we conclude that SpCas9 is capable of cleaving both DNA strands when the targeted sequence and the PAM are offset in either direction by up to 2 base pairs when the DNA substrate is supercoiled.

A time-course of supercoiled DNA cleavage showed that canonically-targeted RNPs (S1_T0, S2_T0, and S3_T0 SpCas9) rapidly reached DSB saturation on the timescale of several minutes. In general, shift-PAM-targeted RNPs with a mild shift (S1_T1, S1_T2, and S1_T-1, but not S1_T-2, SpCas9) displayed similar kinetics of DSB formation as canonically targeted RNPs, whereas shift-PAM RNPs with larger shifts (S1_T3 and S1_T4, SpCas9) slowly accumulated DSBs over three hours (Extended Data Fig.1e).

### The DSB generating activity of shift-PAM-targeted RNPs depends on the canonical PAM

We reasoned that if the shift-PAM mechanism relies on the nearby canonical PAM, mutation of the ‘NGG’ PAM will eliminate the DSB cleavage activity of shift-PAM-targeted RNPs. Indeed, mutating both guanines of the PAM in the S1 locus abolished the DSB generating activity of all canonically- and shift-PAM-targeted RNPs in both supercoiled and linearized DNA substrates (Extended Data Fig.2a-c). While the same result was achieved for S5_T2, S4_T2 required additional mutations to remove alternative PAMs (Hirano et al., 2016) to abolish the DSB cleavage completely (Extended Data Fig.2d,e). We next addressed whether changes within the intervening 2nt offset sequence resulting from use of the S1_T2 crRNA influenced DNA processing. Mutating the first and second adenines adjacent to the canonical PAM to T, G, or C, respectively, did not affect DSB cleavage by S1_T2 SpCas9 (Extended Data Fig.2f, g). Thus, shift-PAM targeting strictly requires the contribution of the canonical PAM and places additional strain on the non-complementary strand.

### Plasticity of endonuclease activities for shift-PAM-targeted complexes

Since single-strand breaks accumulated in supercoiled DNA substrates with 3-5 bp PAM spacing, this suggested the possibility that either the HNH or RuvC domain may be solely responsible for generating such breaks. To test this, we employed two widely-used nickases, D10A SpCas9 possessing a catalytically-active HNH domain but a dead RuvC domain (denoted HNH-SpCas9), and H840A SpCas9 possessing a catalytically-dead HNH domain but an active RuvC domain (denoted RuvC-SpCas9). HNH-SpCas9 was unaffected in its ability to robustly nick substrate as spacing initially increased (S1_T0, S1_T1, S1_T2), but this activity dropped roughly threefold at S1_T3 spacing and decayed to near-zero with S1_T4 spacing (Extended Data Fig. 3a-c). Spacing-dependent incision by RuvC-SpCas9 displayed a slightly more complex pattern, with an increase in incision from S1_T0 to S1_T1 spacing but then a progressive decay as spacing increased to S1_T4, at which incision nevertheless remained at ~60% of the level observed for S1_T0 (Extended Data Fig. 3a-c). Overall, we find that incision of supercoiled substrate by the HNH domain is robust up to S1_T2 spacing but slows beyond that, whereas incision by the RuvC domain is more tolerant of increased spacing over the same scale. Comparing the pattern of incision for increased spacing observed in the two mutants to that observed in wildtype SpCas9 (Figure 1b), we note that incision of supercoiled substrate by wildtype enzyme also significantly dropped threefold upon increasing from S1_T2 to S1_T3, pointing to a dominant role for HNH incision in the response of DNA processing to increased spacing and its enhancement by DNA supercoiling (Figure 1d and Extended Data Fig.1a, c).

### Shift-PAM-targeted SpCas9 generate DSBs in living cells

To determine whether spaced SpCas9 cleavage can be detected in human cells, we first applied it to knock out GFP in K562-iCas9-GFP cells. First we compared the knockout efficiencies of canonical (GFP_T0) and shift-PAM (GFP_T1, GFP_T2) guides, using a generic canonical target in the RAG1 locus (RAG1L_T0) as a negative control. GFP_T1 sustained slightly weaker knockout efficiency (~45%) than GFP_T0 (~55%), while GFP_T2 generated marginal knockout efficiency (Extended Data Fig. 4a-b). Second, we employed high-throughput genome-wide translocation sequencing (LAM-HTGTS) (Hu et al., 2016) in HEK 293T cells to compare translocations to three canonically-targeted (T0) SpCas9 prey DSBs in the RAG1 locus (RAG1B, RAG1C and RAG1D) (Frock et al., 2015) and their shift-PAM-targeted variants (T1, T2) using two independent and canonically-targeted bait DSBs (bait-A and bait-L) located within 1kb of the SpCas9 prey DSBs (Figure 1e-f). From both baits, we recovered several hundreds of translocations from RAG1C_T1 and RAG1D_T1, but not RAG1B_T1, prey sites that were at a similar or modestly reduced level compared to translocations recovered from RAG1C_T0 and RAG1D_T0 prey sites (Figure 1e-f, Extended Table 7). Notably, the translocation peaks for the complementary strand breakpoints were accordingly shifted by the T0 versus T1 targeted SpCas9 prey DSBs (Figure 1e-f). While detected less frequently than T1 DSBs, only RAG1D_T2 DSBs were detected and translocation peaks on the complementary strand correspondingly displayed a 2 bp shift (Figure 1f). Five additional canonically- and shift-PAM-targeted sites (RAG1K-O), using RAG1D_T0 as bait (Bait-D), displayed similar T0 versus T1 profiles, with only RAG1L_T2 forming substantial translocations (Extended Data Fig. 5a, Extended Table 7). Bait/prey LAM-HTGTS experiments generally expressed comparable levels of SpCas9 protein across all conditions (Extended Data Fig. 5b). Therefore, T1 and T2 shift-PAM targeted DSBs can form in cells at levels lower than their canonically-targeted counterparts, yet robustly enough to be detected by translocation.

### Shift-PAM-targeted SpCas9 cleavage leads to 3’ overhangs

SpCas9 DSB products are described predominantly as blunt-ended (Gasiunas et al., 2012; Jinek et al., 2012) but are also capable of generating short 5’ overhangs in cells (Lemos et al., 2018; Liang et al., 2021) that are likely due to increasing cleavage position variance on the non-complementary strand over time (Jinek et al., 2012). However, it is unclear if the cleaved products using shift-PAM spaced sites would predominantly bear blunt ends or overhangs. To avoid the extensive end processing present in cells, we used a combination of two Sanger sequencing approaches, “In-Del” and “5’-locating” (Extended Data Fig. 6a), to obtain the likelihood of end structures from RNPs with shift-PAM guides in biochemistry assays (Extended Data Fig. 6b-j, Extended Table 1). The “In-Del” method is designed to capture the insertion, del etion or direct joining of each targeted DSB, and the “5’-locating” takes advantage of an upstream SbfI site to identify the 5’ ends generated by the SpCas9 RNP (Extended Data Fig. 6a).

Using these sequencing approaches, the near-totality of incision sites on the complementary strand were identified at position 17 from the 5’ end of the 20 nt targeting sequence in all instances assayed (Extended Data Fig. 6b-j; Extended Table 1), faithfully positioned within the RNA:DNA hybrid even as the hybrid was shifted away from the PAM in base-pair increments. The incision site on the non-complementary strand also remained faithfully positioned with respect to the PAM site for up to 2 bp shifts (eg for S1_T0, S2_T0, S1_T1, S1_T2, and S5_T2), while more than half of the observed incision sites for S4_T2 shift by 1bp. However, S1_T3 RuvC-SpCas9 incision was promiscuously distributed across the 3bp spacer region and also in alignment with the HNH cleavage site, whereas S1_T4 RuvC strand cleavage was nearly entirely in alignment with the HNH cleavage position (Extended Data Fig. 6b-i; Extended Table 1). As a result, while S1_T0 and S2_T0 targets displayed blunt ends; S1_T1, S1_T2, S4_T2 and S5_T2 respectively displayed a majority of 1-2 nt 3’ overhangs. S1_T3 generated a mix of overhangs and blunt ends, and S1_T4 generated blunt ends.

We thus also asked whether shifting the RNA:DNA hybrid towards the PAM produced 5’ overhangs however, S1_T-1 targeting resulted nearly entirely in blunt ends that were now 2bp away from the PAM (Extended Data Fig. 6j), indicating steric hindrance by HNH targeting. The altered positional cleavage of the non-template strand in the contexts of S1_T3, S1_T4, and S1_T-1 shows that RuvC endonuclease activity does not require consensus spacing to the NGG sequence recognized by the PID for incision. These combined observations suggest shift-PAM-targeted SpCas9 cleavage relies on a DNA-mediated induced fit mechanism.

### SpCas9 RNP dynamics on single supercoiled DNA molecules

To kinetically characterize the succession of DNA processing steps by shift-PAM targeting RNPs and compare them to processing by canonically-targeted RNPs, we carried out single-molecule assays based on the magnetic trap. Succinctly, single DNA molecules containing target sequences and bearing biotin labels at one end and digoxigenin labels at the other end are attached first to streptavidin-coated magnetic beads and then deposited on and attached to an anti-dig-coated glass slide (Strick et al., 1996). The slide is placed under a microscope and two magnets are used to apply a force on and rotate the magnetic bead. Imaging of the bead position above the surface allows one to measure DNA extension and thus conformation in real-time. For a given, low, fixed force (F = 0.2 pN for the measurements herein), the extension of torsionally-relaxed DNA represents a maximum. Supercoiling the DNA causes it to compact, which is observed in the magnetic trap as a decrease in the “altitude” of the bead above the surface. DNA incision results in an abrupt loss of supercoils, without which the DNA is no longer compact and the bead recovers its position in the torsionally-relaxed state. DNA molecules which do not compact upon supercoiling during the initial calibration steps of the assay are pre-nicked and are excluded from analysis.

Because the torque from positive supercoiling inhibits R-loop formation, RNP injection took place on positively supercoiled DNA extended by 1 pN force. After RNP injection the force was reduced to 0.2 pN and eight negative supercoils were applied to the DNA to initiate single-molecule detection of SpCas9 RNP dynamics: R-loop formation (apparent as DNA unwinding (Ivanov et al., 2020)), single-strand DNA incision (apparent as a sudden loss of supercoiling (Mardenborough et al., 2019)), and double-strand DNA breakage (apparent as terminal loss of the magnetic bead (van den Broek et al., 2005)), see Figure 2e,h and Extended Data Fig. 7b, d; states vi, vii and viii. Using this assay, we could specify the waiting time required to form R-loops (<t_R-on_>, the dwelltime in state vi), the time elapsed from R-loop formation to supercoil dissipation (<t_Super-off_>, the dwelltime in state vii), and the time elapsed from supercoil dissipation to terminal loss of magnetic bead (<t_Bead-off_>, the dwelltime in state viii).

**Figure 2.**
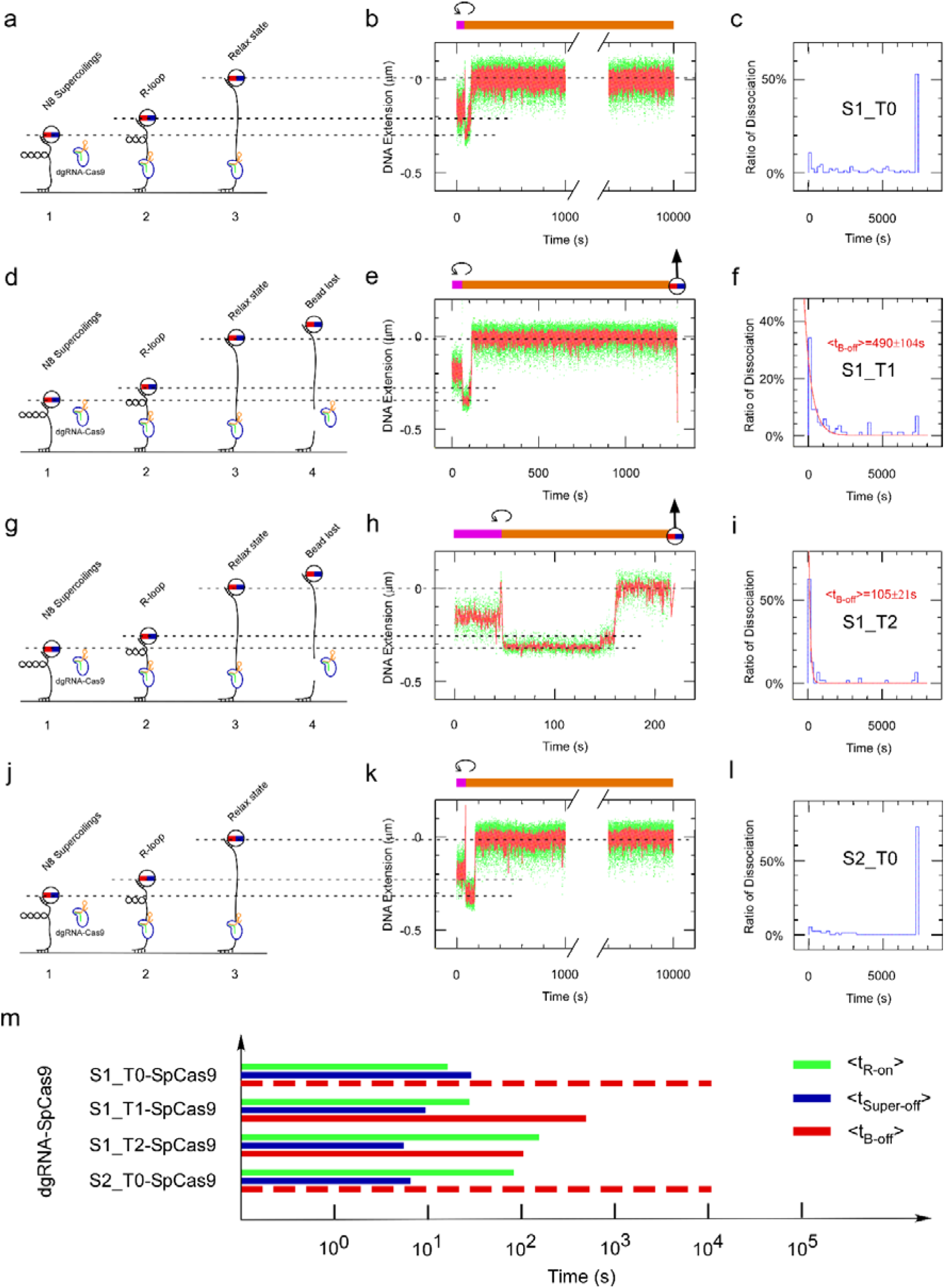
Real-time observation of pre-catalytic and post-catalytic steps by SpCas9 coupled with canonical and shift-PAM guides. **(a-c)** Experiment carried out using S1_T0 SpCas9 RNPs. **(a)** Sketch of assay. SpCas9 RNPs is introduced under non-permissive conditions (positive supercoiling), and (1) its activity initiated by generating 8 negative supercoils in the DNA (see Methods and Extended Data Fig. 7). (2) R-loop formation is detected as an increase in DNA extension and is followed by (3) loss of DNA supercoiling and ultimately (4) loss of the magnetic bead. **(b)** Typical time-trace for the assay. The magenta bar highlights the phase during which DNA is positively supercoiled for the purposes of RNPs injection, the counter-clockwise symbol denotes the moment at which the DNA is negatively supercoiled to initiate the reaction, and the orange bar highlights the phase during which DNA is negatively supercoiled. The magnetic bead with a vertical arrow represents irreversible loss of the magnetic bead. Dash lines are guides to the eye highlighting the bead position in the different states including initial negative supercoiling, R-loop state, and torsionally relaxed state. **(c)** The fraction of molecules forming an R-loop which go on to display terminal bead loss (monitored over 7200 seconds). Following rows are as with the first row but with **(d-f)** S1_T1 SpCas9, **(g-i)** S1_T2 SpCas9, and **(j-l)** S2_T0 SpCas9. **(m)** The timeline summary of events after initiating RNPs activity is displayed in log-scale: R-loop formation (<t_R-on_>, green lines), supercoiling dissociation (<t_Super-off_>, blue lines) and irreversible loss of magnetic bead (<t_B-off_>, red lines (quantified) and dotted red lines (estimated). See Extended Table 2 for detailed measures.

### Unstable shift-PAM-targeted SpCas9 post-cleavage complexes

The first key observation from these measurements is that progressive increases in distance from S1_T0 to S1_T1 and then S1_T2 results in a progressive increase in <t_R-on_> and a progressive decrease in <t_Super-off_> (Figure 2b,e,h,m, Extended Data Fig.8a-c, e-g, Extended Table 2). The increase in <t_R-on_> is consistent with the notion that increased mechanical stress in shift-PAM SpCas9-DNA complexes increases the time required to form the R-loop state. The decrease in <t_Super-off_> suggests that mechanically stressed complexes lead to faster dissipation of supercoils. This could result, somewhat counterintuitively, from faster kinetics for single-strand incision, or alternatively from a reduced ability of stressed complexes to “grasp” the DNA on either side of the nick and prevent DNA from swivelling freely to dissipate supercoils.

Consistent with this alternative explanation, the second key observation is that for canonical S1_T0 and S2_T0 SpCas9 RNPs, <t_Bead-off_> was often too long to measure, typically greater than 7000 seconds (Figure 2c, l). In stark contrast, S1_T1 and S1_T2 complexes resulted in terminal bead loss in only hundreds of seconds (Figure 2f, i). This apparent higher efficiency of DSB induction using S1_T1 and S1_T2 SpCas9 compared to S1_T0 SpCas9 and S2_T0 SpCas9 RNPs is also somewhat counterintuitive given the bulk assay (Figure 1b-d; Extended Data Fig. 9a) which displayed comparable levels of DSBs. To monitor the efficiency of this reaction we calculated the fraction of molecules which progress from R-loop formation to apparent DSB induction by dividing the number of bead-loss events over a two-hour period by the total number of molecules that being able to form R-loops. We found that with this metric, bead-loss fractions were still significantly higher in S1_T1 and S1_T2 than in S1_T0 and S2_T0 (Extended Data Fig. 9b). Analysis carried out on another site (S5, see Extended Data Fig.10a-i) confirmed that the canonical PAM-mediated post-catalytic complex is starkly more stable than the shift-PAM variation. These findings lead us to predict that SpCas9 RNPs confirmed to have incised DNA (i.e. observed to be in the torsionally relaxed state), and in particular canonical SpCas9 complexes which result in relatively fewer lost beads, could perhaps harbour double-strand breaks held together by the SpCas9 RNP.

To test for this, we took advantage of the fact that S2_T0 and S3_T0 SpCas9 target the template strand used by RNA polymerase to transcribe the nanomanipulated DNA. We reasoned that if dissociation of the post-cleavage complex is the rate-limiting step in processing a target via canonical PAM usage, then transcription by RNA polymerase may result in collision with the post-cleavage complex, facilitating its dissociation. Indeed, transcribing RNAP almost doubled the fraction of beads lost (Figure 3a-f; Extended Data Fig. 9a-b). We thus propose that the low rate of bead loss from canonical SpCas9 targeting is due to the relatively higher stability of the post-cleavage DNA-dgRNA-SpCas9 complex compared to shift-PAM targeted SpCas9.

**Figure 3.**
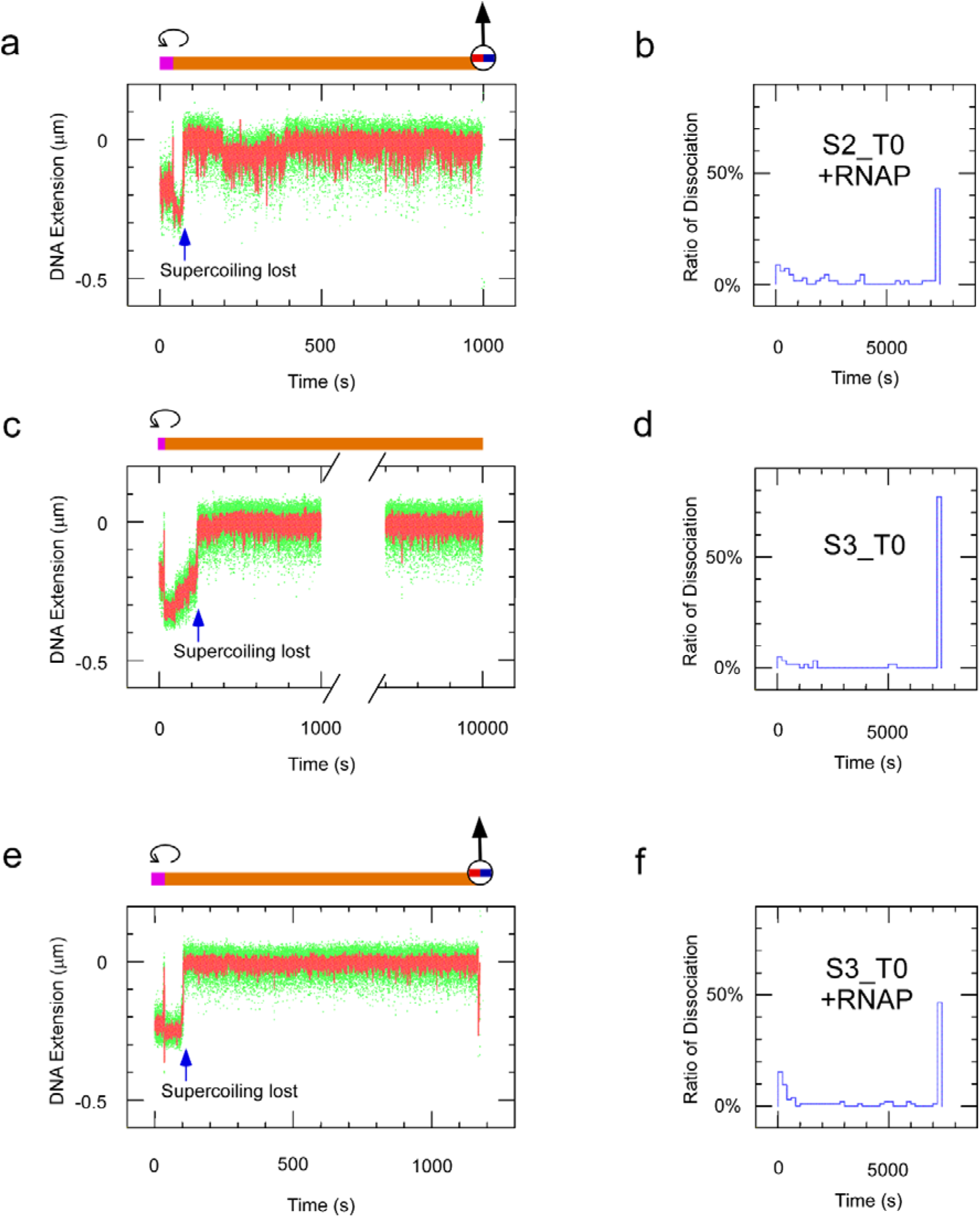
Observation and quantification of terminal bead loss for targeted DNA regions with canonical PAM spacing in the absence or presence of RNAP. **(a)** Typical DNA extension time-trace was obtained using S2_T0 SpCas9 combined with RNAP. DNA was initially positively supercoiled by +5 turns to allow reagent introduction before being unwound by 8 negative turns (counterclockwise arrow) to permit the sequence of events including R-loop formation, supercoil loss, and terminal bead loss (as per Extended Data Fig. 7b). **(b)** The percentage of bead-loss events as normalized by the number of molecules that formed R-loops (as per Extended Data Fig. 7b) shows a bimodal distribution. **(c,d)** and **(e,f)** as for (a,b) but for S3_T0 SpCas9 with and without RNAP, respectively. The mean of bead-dissociation ratio observed within a 2h window for S2_T0 SpCas9 plus RNAPs, S3_T0 SpCas9, and S3_T0 SpCas9 plus RNAP are characterized as 56.5% (n=69), 22% (n=59) and 52.4% (n=103) respectively. All events lasting more than 7200 seconds are transformed as 7200 s.

### HNH domain stability protects incised DNA from rapid loss of supercoiling

Similarly, we propose that faster supercoil dissipation observed in stressed vs. unstressed SpCas9 complexes is due to the relatively higher stability of canonically spaced, unstressed SpCas9 complexes. To explore this hypothesis, we analysed supercoil-loss times for unstressed and stressed HNH-SpCas9 and RuvC-SpCas9 described earlier (Qi et al., 2013).

Results obtained using HNH-SpCas9 hew closely to those obtained with wt-SpCas9, with PAM-shifting resulting in an increase in time required to form the R-loop and a decrease in time required to observe loss of supercoiling (Fig. 4). PAM-shifting with RuvC-SpCas9 also resulted in an increase in time required to form the R-loop and a decrease in time required to observe loss of supercoiling, although it is interesting to note that with this variant a significant fraction of molecules displayed rapid loss of supercoiling even with the non-shifted PAM (Fig. 5 and Ext. Table 2). [We note that differences in the rates of R-loop formation between wild-type and variant SpCas9 should not be interpreted as they may simply be attributed to differences in specific activities as the enzyme preparations are distinct.]

**Figure 4.**
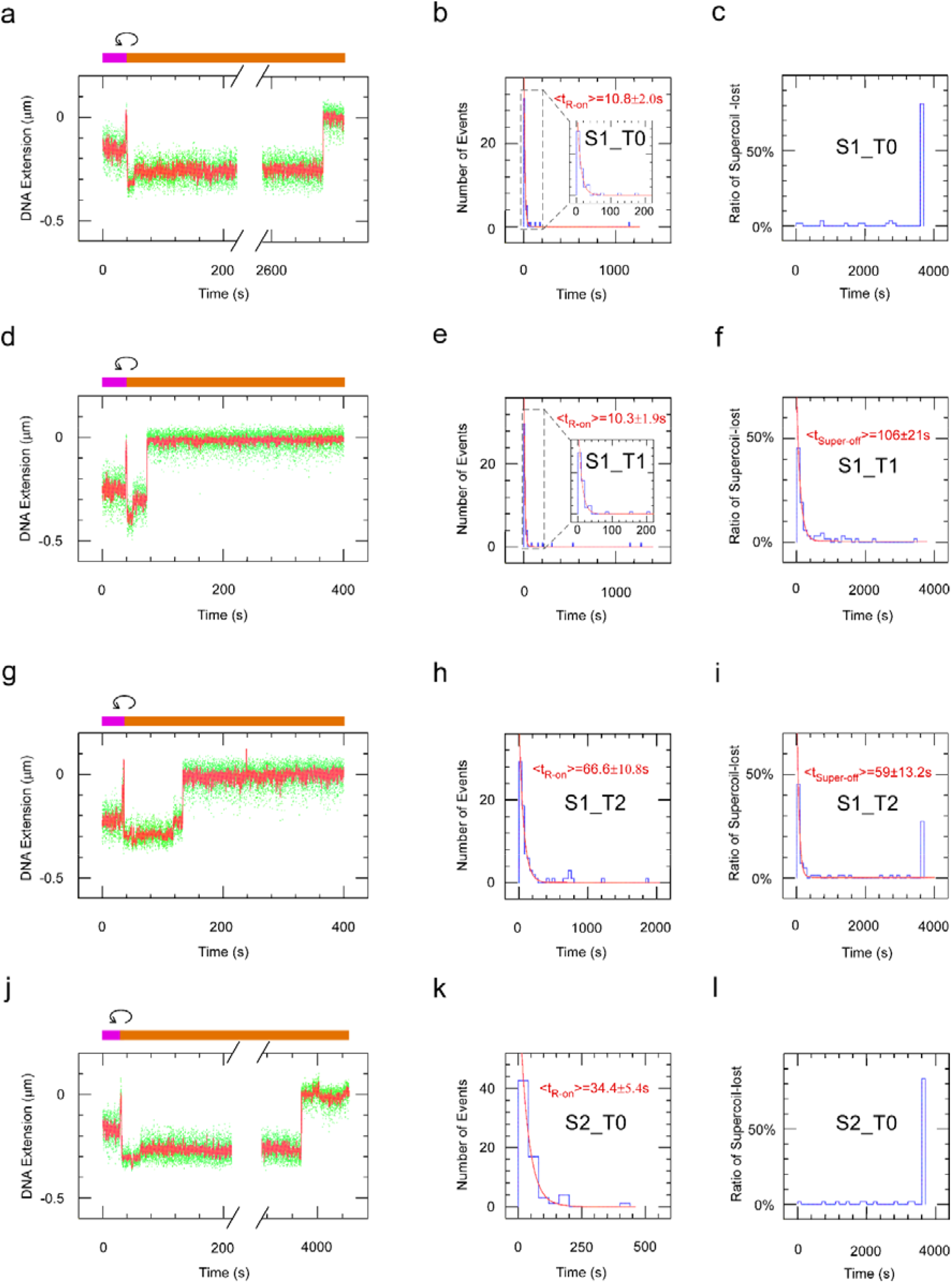
R-loop formation and supercoil loss in DNA nicking assays using HNH-SpCas9 (D10A). **(a-c)** Experiment carried out using S1_T0 HNH-SpCas9. (Left panel) Bead position time-trace. The magenta bar highlights the stage that DNA is positively supercoiled for the purposes of RNPs injection, the counter-clockwise symbol denotes the moment at which the DNA is negatively supercoiled to initiate the reaction, and the orange bar highlights the stage that DNA is negatively supercoiled. The moment of R-loop formation is indicated by a vertical blue arrow, and the moment of supercoil loss is indicated by a vertical red arrow. (Middle panel) Distribution of times for R-loop formation (blue) is fit to a single-exponential distribution (red line), yielding the indicated lifetime of mean. (Right panel) Distribution of time elapsed between R-loop formation and loss of supercoiling (blue) is also fit to a single-exponential distribution (red line) with indicated lifetime of mean. Following panels are as with first three panels but with **(d-f)** S1_T1 HNH-SpCas9, **(g-i)** S1_T2 HNH-SpCas9 and **(j-l)** S2_T0 HNH-SpCas9. For S1_T1 HNH-SpCas9 and S1_T2 HNH-SpCas9, the first modal, i.e., the fast-dissociated supercoil-loss events, are fit to single-exponential distributions (red line) with indicated lifetime of means (**f,i**). All events lasting more than 3600 seconds are transformed as 3600 s. The population of molecules that last > 3600 s are 81.4%, 0%, 28% and 84.1% for S1_T0 HNH-SpCas9, S1_T1 HNH-SpCas9, S1_T2 HNH-SpCas9 and S2_T0 HNH-SpCas9, respectively. See Extended Table 2 for specific measures.

**Figure 5.**
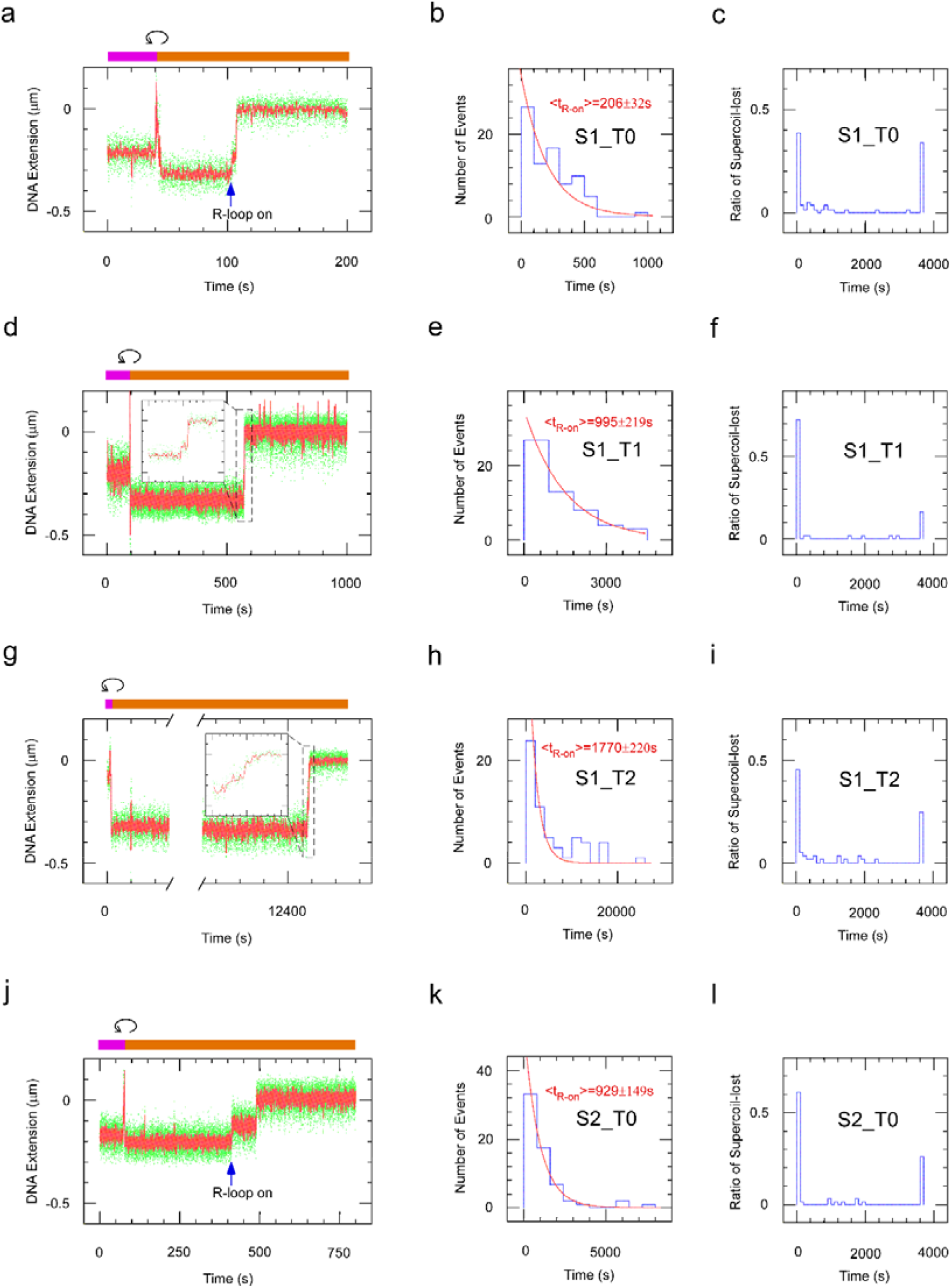
R-loop formation and supercoil loss in DNA nicking assays using RuvC-SpCas9 (H840A). The panels are as with three panel per row in figure 4 but with **(a-c)** S1_T0 RuvC-SpCas9, **(d-f)** S1_T1 RuvC-SpCas9 and **(g-i)** S2_T0 RuvC-SpCas9. The steps of R-loop formation and supercoil loss of the representative time-trace of S1_T1 RuvC-SpCas9 catalytic events (**d**) are shown in the enlarged inner-plot (spanning 560-580s). All events lasting more than 3600 seconds are transformed as 3600 s. The population of molecules that last > 3600 s are 34.6%, 16.4% and 26.2% for S1_T0 RuvC-SpCas9, S1_T1 RuvC-SpCas9 and S2_T0 RuvC-SpCas9, respectively. See Extended Table 2 for specific measures.

The apparent higher efficiency of supercoil-loss events by strained as compared to unstrained HNH-SpCas9 contrasts with the real comparable incision efficiencies seen in ensemble assays (using SDS/heat denaturation). This suggests that HNH-SpCas9 helps to maintain supercoiling despite strand cleavage, presumably because of the RNA “splint,” and that straining the RNP complex with shift-PAM spacing causes it to rapidly lose its grip on the incised yet still supercoiled DNA.

Combined, these observations indicate that a common mechanism – partial RNP dissociation following incision – underlies the observation of supercoil-loss, and that RuvC-SpCas9 activity more readily leads to supercoil loss because of the absence of the RNA splint on the incised strand. We note that supercoil loss for wildtype complexes is slower than observed for RuvC-SpCas9 but faster than observed for HNH-SpCas9, indicating that both stochastic activities contribute to supercoil loss in wildtype complexes and supporting the view that the two incision events are not strictly ordered. Increases in the rate of supercoil loss for wildtype complexes as they are progressively strained from T0 to T2 track increases in the rate of supercoil loss observed for strained HNH-SpCas9, underscoring the rate-limiting role of HNH activity. These results point to destabilization and disassembly, via conformational distortion of the post-catalytic complex at the level of the R-loop itself, as a key rate-limiting feature of the complete processing reaction.

## Discussion

In this study we characterize shifted PAM-mediated DNA processing by SpCas9. We propose that when NGG PAMs are not canonically positioned adjacent to the RNA-guided target sequence, SpCas9 retains the ability to “scrunch” several nucleotides of substrate (Revyakin et al., 2006), achieving double-strand DNA cleavage by off-set incision of the two strands and destabilizing the post-cleavage complex. Consistent with this “scrunching” hypothesis for R-loop formation using shifted PAMs, we compared the size of the R-loop generated by canonical (S1_T0) and strained (S1_T2) SpCas9-DNA complexes and found that the R-loops generated by the latter are as predicted ~2bp larger than those generated by the former (Figure 6a-b) (Strick et al. 1998, Szczelkun et al., 2014).

**Figure 6.**
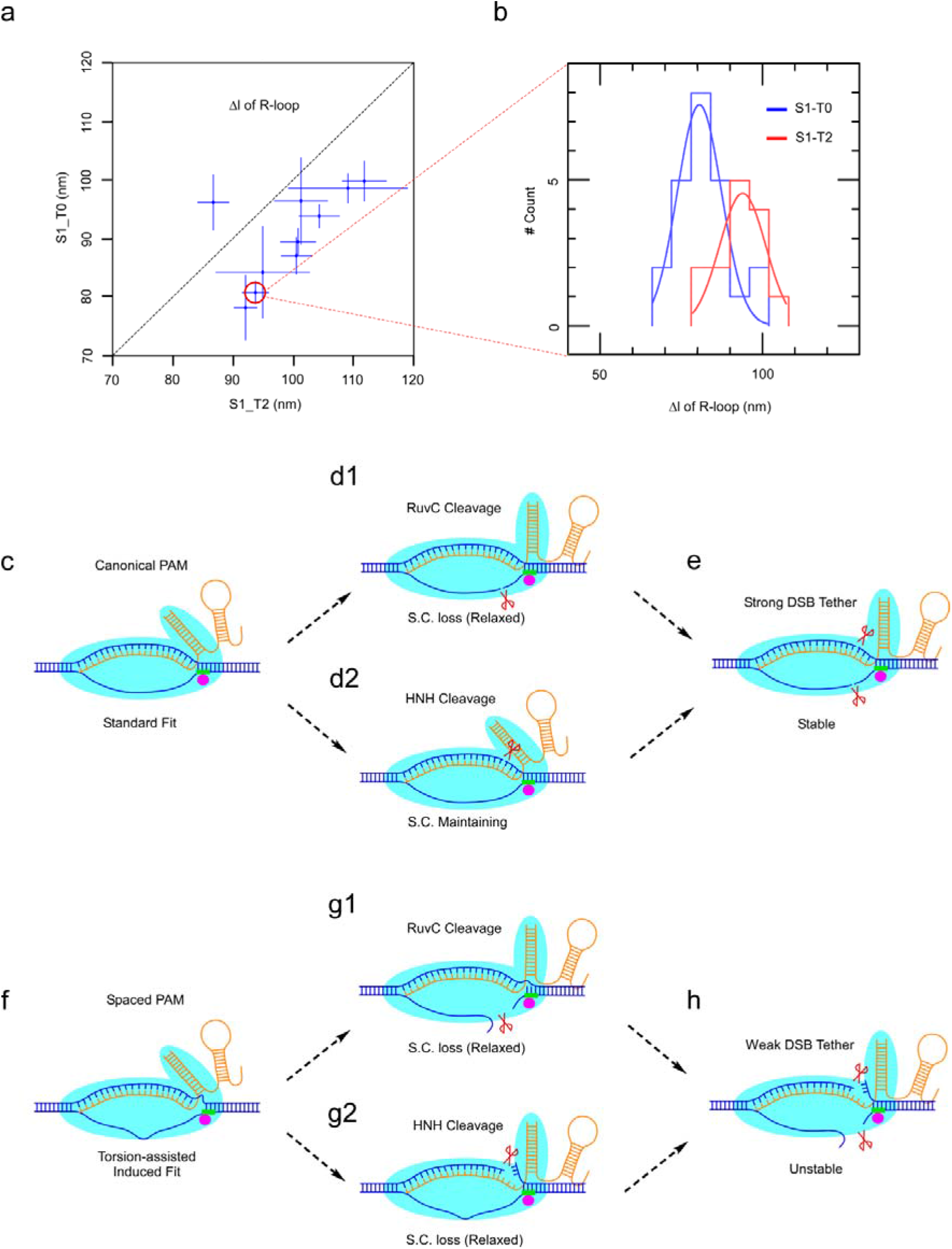
DNA scrunching in shift-PAM complexes and model for catalytic steps on target sequences with canonical and strained PAMs. **(a)** Negatively supercoiled DNA molecules were treated with S1_T2 dSpCas9 to generate an R-loop and determine the corresponding increase in DNA extension, after which S1_T2 dSpCas9 RNPs were washed out and the same DNA molecules were then treated with S1_T0 dSpCas9 to again generate an R-loop and determine the corresponding increase in DNA extension. Mean extension changes observed on a given DNA molecule as resulting from the two different RNPs are shown in a square with a 45-degree diagonal (dashed line), along with the associated error bars (SEM). Complexes were repeatedly formed on negatively supercoiled DNA and ejected by positively supercoiling the DNA so as to accumulate statistics on each DNA molecule. **(b)** Histograms of observed DNA extension changes upon exposure to the two different RNPs above are shown for a representative DNA molecule and are fit to Gaussian distributions. The average increase in DNA extension upon R-loop formation with S1_T0 and S1_T2 dSpCas9 RNPs was determined as 81 ± 2 nm (SEM, n = 26 events) and 94 ± 3 nm (SEM, n=19 events), respectively, corresponding to an extra ~2 bp of unwinding for the S1_T2 dSpCas9-based R-loops (see Methods). A minor fraction (15%) of presumably immature R-loops with only about ½ the full unwinding (only observed using dSpCas9) were excluded from analysis. **(c-e**) Model for catalytic steps on target sequence with canonical PAMs: **(c)** the first 20 nts of crRNA forms an R-loop with the complementary strand, and the PAM interacts with the PAM-interacting motif of SpCas9. **(d1, d2)** The complementary and non-complementary strand can be incised by either the HNH nuclease or the RuvC nuclease, respectively. Incision of the complementary strand by the HNH nuclease does not favour loss of supercoiling because the RNA:DNA hybrid acts as a splint across the cut site. In contrast, incision of the non-complementary strand by the RuvC domain tends to result in loss of supercoiling. **(e)** Upon cleavage of both strands, the stable post-cleavage complex can tether the broken DNA ends for long times. **(f-h)** In the pre-catalytic complex formed with a shifted PAM, **(f)** mechanical strain slows down R-loop formation and PAM binding. **(g1, g2)** However, mechanical strain also results in the instability of such complex, with signature loss of supercoiling upon incision by the HNH domain as well as the RuvC domain. **(f)** Once double-strand cleavage is completed, the strained, destabilized post-cleavage complex holds the broken DNA ends together more weakly, favoring terminal separation of DSB ends. DNA is in blue, crRNA tracrRNA duplex is in orange, PAM is in green, cleavage sites are indicted by scissors, SpCas9 is in cyan rabbit-like shape, and the PAM-interacting motif is in magenta dot.

Using bulk assays and sequencing assays, we show that strained SpCas9 complexes generate DNA overhangs of up to a few bases in length. As strain increases, HNH domain incision of the complementary strand faithfully tracks the RNA:DNA hybrid position as it moves away from the PAM, maintaining incision 17 nt from the 5’ end of the hybrid. RuvC incision remains adjacent to the PAM for up to two bases of strain, resulting in robust generation of up two-base 3’ overhang at the DNA ends. However beginning at three bases of strain RuvC domain incision is seen to “slip” from PAM-adjacent to HNH-adjacent, presumably highlighting the distinct sets of interactions between the RuvC domain and the HNH domain and the RuvC domain and the PID. The RuvC domain can still incise the non-complementary strand in the “slipped” state but these incision events face the HNH incision site and once again result in blunt overhang formation. In human cells, the 1 bp-shifted PAMs lead to robust DSBs, however, 2bp-shifted PAMs generate much weaker cleavage probably due to the less negatively-supercoiled nucleosome-bound genomic DNA in cells.

Single-molecule interrogation of the kinetics of R-loop formation, DNA single-strand incision, and DNA double-strand cleavage by SpCas9 and its nuclease domain mutants in the context of canonical and shift-PAM targeting provides us a deeper understanding of SpCas9 cleavage and how the pre-catalytic and post-catalytic complexes are affected by offsetting the guide/PAM spacing. This is consistent with earlier findings that the DNA bubble within the seeding region can tolerate RNA:DNA mismatches (Sternberg et al., 2015). Taken together, these results lead us to propose a unified catalytic mechanism of SpCas9 action which is modulated by the spacing between the PAM and the R-loop. For canonically-spaced PAMs (Figure 6c-e), PID binding contributes to stabilizing the pre-catalytic complex (Figure 6c), and is followed by complementary strand cleavage by the HNH domain (Figure 6d1) or non-complementary strand cleavage by the RuvC domain (Figure 6d2) in a stochastic manner. While HNH nuclease incision within the R-loop does not lead to immediate loss of supercoils, presumably because the RNA can act as a “splint”, RuvC nuclease incision in the non-complementary strand leads to rapid loss of supercoils. Once both strands are incised, the two broken DNA ends remain tethered thanks to the overall stability of the complex bridging the break (Figure 6e). In the case of shifted PAMs additional mechanical stress is introduced in the R-loop state, overcoming the stabilizing effect of the splint and resulting in rapid loss of supercoils even upon HNH-domain incision and rapid dissociation of the post-catalytic complex once both strands have been cleaved (Fig. 6f-h).

Consistent with our model, recent studies determined that the complementary strand is more stable than the non-complementary strand in the post-catalytic complex (Wang et al, 2021; Richardson et al., 2016). From the single-molecule and bulk assays carried out here, we conclude that SpCas9-mediated double-strand cleavage of DNA does not necessarily lead to rapid dissociation of broken ends. Supercoiling is seen to be rapidly lost in unstrained wtCas9, with both RuvC and HNH activities participating in the first incision event. The rate of supercoiling loss increases with strain. For unstrained SpCas9 in just over half of all events secondary incision appears to be hidden from detection by SpCas9 remaining on the DNA break, most likely bridging the break by maintaining the R-loop and the PID engaged in the post-incision complex. The complex can be destabilized by strain in the spacing between the RNA:DNA hybrid and the PAM, leading to a much higher rate of terminal bead loss for strained complexes. The complex can also be driven off using transcribing RNA polymerase. Strained complexes thus result in enhanced disassembly from DNA, which here appears as a significant yet overlooked rate-limiting step to DNA processing.

Our work complements and extends previous studies demonstrating that SpCas9 is much less tolerant of guide DNA mismatches in the PAM-proximal seeding region than those occurring in PAM-distal non-seed regions (Yang et al., 2017), and that directed mutations in SpCas9’s PAM-interacting domain can result in variants preferring non-canonical PAM combinations (Kleinstiver et al., 2015; Hirano et al., 2016; Anders et al., 2016). The shift-PAM targeting described here further expands options when designing sgRNAs and will be useful as applied to various genome-editing contexts.

The combination of ensemble assays, next generation sequencing tools and single-molecule studies presented here allow us to dissect the sequence of steps from R-loop formation, to single-strand break induction, to double-strand break formation, to post-incision complex dissociation by SpCas9 acting on its target DNA using both canonical and strained R-loop/PAM spacing. Our results demonstrate that the protein-DNA complex is sufficiently deformable as to tolerate modulation of the distance between PAM and seed region, and that this is sufficient to cause a single mechanically-stressed dgRNA-SpCas9 to generate overhanging ends. This provides an additional degree of freedom in choosing SpCas9 targets, and an added layer to discern potential off-target effects during guide RNA design.

### Limitations of the study

The single molecule assays revealed that post-catalytic complex dissociation is a rate-limiting step for separating DNA ends, and that shifted PAMs speed up dissociation. To directly visualize the SpCas9 dynamics and to address if SpCas9 remains on either of the two ends after end separation, other single molecule tools including TIRF and FRET need to be employed.

SpCas9 can generate short, templated insertions, likely involving staggered cleavage and 5’-overhangs, but the molecular mechanism allowing this remains unclear (Zou et al., 2016; Shou et al., 2018; Shen et al., 2018; Allen et al., 2018; Lemos et al., 2018; Jones et al., 2021; Liang et al., 2021). Living cell cleavage assay combined LAM-HTGTS show both canonical PAM and 1bp strained PAM generate comparable cleavage frequencies, though larger off-sets significantly decrease cleavage efficiency. Thus, more effective shift-PAM engineering would likely be necessary to potentially enhance gene knock-in efficiency.

## Acknowledgements

We thank Professor Marc Nadal for discussion, and Shuang Wang for a generous gift of RNAP. We thank Marie Le Bouteiller and Paul George Barghouth for advice on data analysis. We thank the Ligue Nationale Contre le Cancer and Agence Nationale pour la Recherche (ANR-17-CE11-0042) for providing financial support. R.L.F is a V Scholar for the V Foundation for Cancer Research.

## Author Contributions

J.L.W. and T.R.S. conceived the project. J.L.W. and J.L.G carried out the experiments and data analysis, J.L.W, R.L.F., and T.R.S wrote the paper.

## Methods

### Cell Lines

HEK293T cells were obtained from ATCC and cultured at 37 °C and 5% CO2 in Dulbecco’s Modified Eagle’s Medium (DMEM) supplemented with 10% (vol/vol) FCS, 50 U/mL penicillin/streptomycin, 2 mM L-glutamine, 1× MEM-NEAA, 1 mM sodium pyruvate, 50 μM 2-mercaptoethanol and 20 mM HEPES (pH 7.4). HEK293T cells maintained in 2.5% FCS (vol/vol) DMEM media only in the period of two hours before transfection and six hours post transfection.

K562-iCas9-GFP cell line was established by infecting the K562 cells (originated from ATCC) with lentivirus carrying Doxycycline-inducible SpCas9-T2A-GFP and BSD gene segments. Briefly, the donor plasmid was engineered by replacing the NeoR of Lenti-iCas9-Neo (Cao et al., 2016, Addgene #85400) with BSD. Infected K562 cells were seed into 96-well at concentration of 0.3 cell/ well. Cell colonies were picked up 10 days later, tested by genotyping, fluorescent microscopy, flow-cytometry and western blot to choose a stable K562-iCas9-GFP cell line. The K562-iCas9-GFP cells were maintained in Roswell Park Memorial Institute (RPMI)-1640 medium supplemented with 10% (vol/vol) FCS, 50 U/mL penicillin/streptomycin, 2 mM L-glutamine, 1× MEM-NEAA, 1 mM sodium pyruvate, 50 μM 2-mercaptoethanol, 20 mM HEPES (pH 7.4) and 10 μg/ml Blasticidin. 1 μg/ml Doxycycline was added only when Cas9 required to be expressed. Doxycycline-treated cells were not used for purposes of cell line preservation.

### RNP- and RNAP-particles Reconstitution in vitro

The crRNAs and tracRNA (see Extended Table 3) were mixed 1:1 to 1 μM in supplied buffer 3.1 (NEB) with 0.2 U/μl RNaseOUT™ (Invitrogen), denatured at 90°C for 2 mins and cooled down at room temperature for 30 mins to allow the formation of RNA duplex dgRNA. SpCas9 and its mutants were added into the dgRNA dimer as 1:1 ratio and incubated at room temperature for another 30 mins to allow forming stable RNP particles.

E.coli RNAP core subunits and σ factor were expressed, purified and reconstituted into holo-RNAP enzyme as described elsewhere (Fan et al., 2016).

### DNA Substrates Preparation

The targeting DNA is a transcript unit of about 200 bps (shown in Extended figure 1), which was inserted into the unique Kpn1 site of the Charomid-9-5 vector. The targeting DNA transcript with ~1 kb arms flanking in each site was amplified by PCR to obtain 2.2 kb DNA inserts and sub-cloned insert into SbfI and XbaI sites of pUC18 vector (Graves et al., 2015). The pUC18 bearing the 2.2 kb DNA inserts and its SbfI-linearized products were the substrates used for ensemble assays. The site-directed mutants were generated by paired PCR and confirmed by Sanger sequencing.

The pUC18 plasmids bearing 2.2 kb targeting DNA were transformed, amplified using E.coli Turbo (NEB), and extracted by NucleoBond midi-prep (Macherey Nagel). The plasmid was further purified by 1% agarose gel electrophoresis and NucleoSpin kit (Macherey Nagel) to obtain supercoiled double-stranded circular plasmids. Linearized DNA substrates were generated by treating the supercoiled plasmids with SbfI, and purified by 1% agarose gel electrophoresis and gel extraction kit.

### Ensemble Biochemistry Assays

Supercoiled and linearized plasmids were mixed with dgRNA SpCas9 or dgRNA SpCas9 mutants as final concentration of 10nM (substrate): 100 nM (RNPs) in 10 ul reaction buffer (20 mM HEPES-K pH 7.8, 150 mM KCl, 5 mM MgCl2, 2 mM DTT, 0.5 mg/ml BSA, 0.05% Tween-20, 0.2 U/μl RNaseOUT™), incubated at 34°C. The reactions were quenched by adding 6x loading dye (with 1% SDS) and incubated at 65°C for 3 mins before running 1% agarose-Etbr gel electrophoresis to detect the cleavage efficiency by ChemiDoc system (Bio-Rad).

### Junction Mapping via Sanger sequencing

“In-Del” assay: the supercoiled plasmids were mixed with dgRNA SpCas9 as final concentration of 10 nM (substrate), 100 nM (RNPs) in 100 μl, and incubated at 37 °C overnight. 6x loading dye was added and incubated at 65 °C for 3 mins, followed by running 1% agarose gel electrophoresis to purify the plasmid with DSBs. The purified DNA molecules were treated with quick-blunting kit (NEB) and ligated using either the quick-ligase kit (NEB) or T4-DNA ligase (NEB) following the corresponding protocols. The re-ligated plasmids were transformed into Turbo competent cells (NEB), incubated on ice for 30 mins, heat-shocked at 42 °C for 45 seconds, cooled down on ice for another 2 mins, briefly recovered in SOD medium at 37 °C for 10 mins and finally spread on LB-agar with Ampicillin and incubated at 37 °C overnight. The next day, single colonies were picked, amplified and plasmid-purified for Sanger sequencing.

“5’-locating” assay: all the procedures are the same as for the “In-Del” assay except the substrate DNA molecules were SbfI-linearized after SpCas9 cleavage. The cleaved products by dgRNA SpCas9 RNPs were about 4 kb and 1 kb, and the 4 kb DNA molecules were subjected to the same blunting, re-ligation, transformation and sequencing as above. Since the 5’ ends of cleaved DNA products would be fixed, the 5’ ends could be determined.

“In-Del” assay is sufficient to obtain single-nucleotide resolution in the case of deletion or insertion of nucleotides in a ATGC-diverse micro-locus (S4_T2 in Extended Table 1); however, it is insufficient to know the precise cleavage site in the cases dominated by perfect repair (S2_T0, S1_T0, S1_T4 and S1_T-1 in Extended Table 1), neither in the cases of insertion or deletion nucleotides in a micro-locus with consecutively identical nucleotides (S1_T1, S1_T2, S1_T3 and S5_2 in table S1). Thus the “5’-locating” assay is required to rule out other cleavage possibilities and to pinpoint the cleavage sites.

### DSB Detection in Living Cells by LAM-HTGTS

HEK293T cells were split at 30% confluence in 6-well plate, cultured in DMEM media with 10% FCS. After 16 hours, the culture media was replaced by low serum media DMEM with 2.5% FCS and maintained for 1-2 hours. Meanwhile, plasmid PEI (Polyethylenimine) mixture was prepared. In brief, 15 ul PEI (1mg/ml) was added into 110 ul NaCl (150 mM) to form PEI working solution, and in parallel, pX330 plasmid (about 1μg/μl each) bearing three T0-sgRNA including RAG1B_T0, RAG1C_T0, RAG1D_T0 (Extended Table 5) were mixed with bait-L or bait-A with ratio 1: 1: 1: 2 (For bait-L and bait-H, 5μg: 5μg: 5μg: 10μg, while for bait-A, 4μg: 4μg: 4μg: 8μg) and added NaCl (150 mM) up to 125 ul to form DNA working solution. The PEI working solution was added into DNA working solution drop by drop and mixed well, incubated at RT for 20 mins and distributed into HEK293T cell culture evenly. The plasmid PEI mixtures for T1 sgRNA, T2 sgRNA and bait-alone control were prepared and distributed into HEK293T cell culture accordingly. After six hours, the low serum media was replaced by growth media DMEM with 10% FCS. After three days, the HEK293T cells were harvested, lysed, and genomic DNA were extracted. Similar practice to extended preys RAG1K-O (3μg each) using RAG1D_T0 as bait (6μg).

For LAM-HTGTS, 50 μg genomic DNA in each treatment condition was prepared into one “library” with unique barcode as described elsewhere (Hu et al., 2016). In brief, the genomic DNA was sheared by sonication to form fragment between 200 bp to 2 kb, followed by LAM-PCR using a biotin-labelled primer close to the bait-site. The products from the LAM-PCR were attached to streptavidin magnetic beads, subjected to adapter-ligation. The ligated products were further undergone nested-PCR using one primer (AP2I7) matched the adapter and another primer (nested primer with several nucleotide barcode) matched region between bait-site and biotin labelled primer, followed by tagged-PCR using primer P7I7 and P5I5 that match the primers used in nested-PCR. The products range between 500 bp to 1 kb from tagged-PCR were purified by 1% agarose gel, subjected to quality control and NovaSeq (Novogene). See Extended Tables 6 for oligos used.

### Western Blotting

The HEK293T cells control and those transfected with various combinations of bait and prey guide RNAs were collected three days post transfection. Five million cells were lysed. Indicated antibodies (anti-Flag, 1:2000, 12793S, Cell Signaling Technology; anti-Rabbit-IgG, 1:2000, G-21234, Thermo Scientific; anti-β-actin, 1:4000, SC-47778, Santa Cruz Biotechnology) and HRP substrate (K-12042-D10, Advansta) were used for the western blotting experiments.

### GFP Knockout and Detection by Flow Cytometry

The gene of sgRNAs GFP_T0, GFP_T1, GFP_T2 and RAG1L_T0 were cloned into pMCB320 plasmid, respectively (Han et al., 2017, Addgene #89356). 5μg GFP_T0-, GFP_T1-, GFP_T2- and RAG1L_T0-pMCB320 plasmids was delivered into K562-iCas9-GFP cells by 4D-nucleofector (Lonza), respectively. Nucleofected cells were recovered in RPMI media with 10% FCS, 0.2mg/ml G418, 5mg/ml Blasticidin and 1mg/ml Doxycycline. The nucleofection efficiencies were measured 2 days post-nucleofection as ~ 60% for all conditions by flow cytometry (BD), taken the advantage of mCherry in pMCB320 plasmid. The efficiency of GFP knockouts were measure on day 7 post-nucleofection by flow cytometry.

### Single-Molecule Manipulation

The 2.2 kb DNA was ligated with 1 kb biotin- and dig-labeled DNA through SbfI- and XbaI-overhangs by T4 DNA ligase (NEB Biolabs). The ligated products were attached to the anti-dig treated surface of reaction chamber and streptavidin-coated magnetic beads, respectively. The attached DNA constructs were the substrates employed for single-molecule assays. All SpCas9 components including dgRNA alone, dgRNA-SpCas9 or dgRNA-SpCas9 mutants were added as 1nM of 150 μl. In transcription-coupled cleavage assay, 200 pM RNAP and 100 μM NTPs were also introduced.

DNA hat-curves were taken at 0.2 pN to distinguish the intact double-strand DNA, nicked DNA and tangled DNAs based on biophysical properties including DNA elasticity and extension (Strick et al., 1996). Only the intact double-strand DNA molecules were employed in following single-molecule experiments. The intact double-strand DNA molecules were re-calibrated at 0.2 pN with 8 turns of negative supercoilings, free state and 5 turns of positive supercoilings for double-check and generating references of DNA extension. Following that, the applied force was increased to 1 pN (the DNA molecules remains with 5 turns of positive supercoilings), and the sample including dgRNA alone, dgRNA-SpCas9 or dgRNA-SpCas9 mutants were introduced in the reaction chamber in each assay. A fluctuation spike caused by the flow of sample injection. When the reaction system become stable again, the force was lowered back to 0.2 pN. The DNA molecules were either kept at 5 turns of positive supercoilings or 8 turns of negative supercoilings to observe the R-loop, supercoilings dissociation and magnetic bead-lost.

### Biochemistry Data Analyses

The cleavage efficiencies of DSBs and SSBs in EtBr-agarose gels were quantified using ImageJ and normalized by subtracting the backgrounds of controls. The efficiencies were further analysed, and corresponding figures were plotted by GraphPad.

### Single-Molecule Data Analysis

Single-molecule data were processed using the PicoJAI software suite (PicoTwist SARL).

#### Temporal analysis

The time required for R-loop formation/dissociation, the time required to transition from the R-loop state to the torsionally-relaxed state (loss of supercoiling), and the time required to transition from the torsionally-relaxed state to irreversible loss of the magnetic bead, were characterized at the level of individual molecular events by analysing DNA extension vs time traces obtained on individual DNA molecules. The mean dwelltimes for the corresponding states observed in pre- and post-catalytic complexes were obtained by fitting the resulting distributions of times to single-exponential decays. Python and GraphPad also used for comparing the significant level of different treatments.

#### Spatial analysis

Nanometer-scale changes in DNA extension during R-loop formation/dissociation were characterized at the level of individual molecular events by analysing DNA extension vs time traces obtained on individual DNA molecules, and then converted to DNA unwinding in base-pairs by converting changes in DNA writhe to changes in DNA unwinding via the linking number formalism as described previously (Strick et al. 1998, Revyakin et al. 2006), using the calibrated change in DNA extension of 55 nm per unit of writhe (in our experimental conditions of force and salinity, see Extended Data Fig. 7a) and a DNA pitch of 10.5 bp/turn. Thus for instance since the seeding sequence of each crRNA is 20 nts, in theory the fully hybrid RNA:DNA (mature R-loop) unwinds approximately two turns of DNA, resulting in the addition of two units of positive writhe into the DNA and a concomitant increase in the extension of negatively supercoiled DNA of about 110 nm. Following single-strand DNA incision, the DNA molecules have the potential to reach the torsionally-relaxed-state (complete loss of supercoiling, corresponding to the maximum-extension state seen in Extended Data Fig. 7a), and following cleavage of both strands, the DNA molecules have the potential to break as seen by irreversible loss of the magnetic bead.

### LAM-HTGTS Data Analyses

LAM-HTGTS data analysis was conducted as previously reported (Hu et al., 2016). In brief, illumina reads were de-multiplexed and adapter sequence trimmed using the fastq-multx tool from ea-utils (http://code.google.com/p/eautils/) and the SeqPrep utility (https://github.com/jstjohn/SeqPrep) respectively. Reads were normalized by Seqtk (https://github.com/lh3/seqtk) and mapped to the hg19 reference genome using TranslocWrapper to map translocations. Reads were also aligned to the bait region using HTGTS Rep to identify rejoining events and then combined to translocations using Rejoin. We note duplicates of translocations were removed to eliminate the artificial duplication of PCR, with features of exact bait sequence and exact prey sequence, which may not be perfect since some of them may be true biological duplications.

## Data Availability

LAM-HTGTS sequencing data are deposited at the Gene Expression Omnibus with the accession number GSE192459. Raw single-molecule time-traces are available upon reasonable request.

## References

Allen F, Crepaldi L, Alsinet C, Strong AJ, Kleshchevnikov V, De Angeli P, et al. Predicting the mutations generated by repair of Cas9-induced double-strand breaks. Nat. Biotechnol. 37: 64–72 (2019).

Anders C, Niewoehner O, Duerst A, Jinek M. Structural basis of PAM-dependent target DNA recognition by the Cas9 endonuclease. Nature 513(7519) 569–573 (2014).

Anders C, Bargsten K, Jinek M. Structural Plasticity of PAM Recognition by Engineered Variants of the RNA-Guided Endonuclease Cas9. Mol. Cell 61(6) 895–902 (2016).

Barrangou R, Fremaux C, Deveau H, Richards M, Boyaval P, et al. CRISPR provides acquired resistance against viruses in prokaryotes. Science 315(5819) 1709–1712 (2007).

Brouns SJ, Jore MM, Lundgren M, Westra ER, Slijkhuis RJ, Snijders AP, et al. Small CRISPR RNAs guide antiviral defense in prokaryotes. Science 321(5891) 960–964 (2008).

Cao J, Wu L, Zhang SM, Lu M, Cheung WK, Cai W, Gale M, Xu Q, Yan Q. An easy and efficient inducible CRISPR/Cas9 platform with improved specificity for multiple gene targeting. Nucleic Acids Res 44(19): e149 (2016).

Cong L, Ran FA, Cox D, Lin S, Barretto R, et al. Multiplex genome engineering using CRISPR/Cas systems. Science 339(6121) 819–823 (2013).

Deltcheva E, Chylinski K, Sharma CM, Gonzales K, Chao Y, Pirzada ZA, et al. CRISPR RNA maturation by trans-encoded small RNA and host factor RNase III. Nature 471(7340) 602–607 (2011).

Fan J, Leroux-Coyau M, Savery NJ, Strick TR. Reconstruction of bacterial transcription-coupled repair at single-molecule resolution. Nature 536(7615) 234–237 (2016).

Frock RL, Hu J, Meyers RM, Ho YJ, Kii E, Alt FW. Genome-wide detection of DNA double-stranded breaks induced by engineered nucleases. Nat. Biotechnol. 33(2) 179–186 (2015).

Garneau JE, Dupuis MÈ, Villion M, Romero DA, Barrangou R, et al. The CRISPR/Cas bacterial immune system cleaves bacteriophage and plasmid DNA. Nature 468(7320) 67–71 (2010).

Gasiunas G, Barrangou R, Horvath P, Siksnys V. Cas9-crRNA ribonucleoprotein complex mediates specific DNA cleavage for adaptive immunity in bacteria. PNAS 109(39) : E2579–2586 (2012).

Graves ET, Duboc C, Fan J, Stransky F, Leroux-Coyau M, Strick TR. A dynamic DNA-repair complex observed by correlative single-molecule nanomanipulation and fluorescence. Nat Struct Mol Biol. 22(6) 452–457 (2015).

Han K, Jeng EE, Hess GT, Morgens DW, Li A, Bassik MC. Synergistic drug combinations for cancer identified in a CRISPR screen for pairwise genetic interactions. Nat. Biotechno 35(5): 463–474 (2017).

Hirano S, Nishimasu H, Ishitani R, Nureki O. Structural Basis for the Altered PAM Specificities of Engineered CRISPR-Cas9. Mol. Cell 61(6) 886–894 (2016).

Hsu PD, Lander ES, Zhang F. Development and applications of CRISPR-Cas9 for genome engineering. Cell 157(6) 1262–1278 (2014).

Hsu PD, Scott DA, Weinstein JA, Ran FA, Konermann S, Agarwala V, et al. DNA targeting specificity of RNA-guided Cas9 nucleases. Nature Biotechnology, 31(9) 827–832 (2013).

Hu J, Meyers R, Dong J. et al. Detecting DNA double-stranded breaks in mammalian genomes by linear amplification–mediated high-throughput genome-wide translocation sequencing. Nat Protoc 11: 853–871 (2016).

Ivanov IE, Wright AV, Cofsky JC, Aris KDP, Doudna JA, Bryant Z. Cas9 interrogates DNA in discrete steps modulated by mismatches and supercoiling. PNAS, 117(11) 5853–5860 (2020).

Jinek M, Chylinski K, Fonfara I, Hauer M, Doudna JA, Charpentier E. A programmable dual-RNA-guided DNA endonuclease in adaptive bacterial immunity. Science 337(6096) 816–821 (2012).

Kleinstiver BP, Prew MS, Tsai SQ, Topkar VV, Nguyen NT, et al. Engineered CRISPR-Cas9 nucleases with altered PAM specificities. Nature 523(7561) 481–485 (2015).

Lemos BR, Kaplan AC, Bae JE, Ferrazzoli AE, Kuo J, Anand RP, Waterman DP, Haber JE. CRISPR/Cas9 cleavages in budding yeast reveal templated insertions and strand-specific insertion/deletion profiles. PNAS, 115 (9) E2040–E2047 (2018).

Liang Z, Kumar V, Le Bouteiller M, Zurita J, Kenrick J, et al. Ku70 suppresses alternative end joining in G1-arrested progenitor B cells. PNAS, 118(21):e2103630118 (2021).

Mardenborough YSN, Nitsenko K, Laffeber C, Duboc C, Sahin E, et al. The unstructured linker arms of MutL enable GATC site incision beyond roadblocks during initiation of DNA mismatch repair. Nucleic Acids Res. 47(22) 11667–11680 (2019).

Marraffini LA, Sontheimer EJ. Self versus non-self discrimination during CRISPR RNA-directed immunity. Nature 463(7280) 568–571 (2010).

Morgens DW, Wainberg M, Boyle EA, Ursu O, Araya CL, et al. Genome-scale measurement of off-target activity using Cas9 toxicity in high-throughput screens. Nat. Commun. 5(8): 15178 (2017).

Revyakin A, Liu C, Ebright RH, Strick TR. Abortive initiation and productive initiation by RNA polymerase involve DNA scrunching. Science 314(5802) 1139–1143 (2006).

Qi LS, Larson MH, Gilbert LA, Doudna JA, Weissman JS, et al. Repurposing CRISPR as an RNA-guided platform for sequence-specific control of gene expression. Cell 152(5) 1173–83 (2013).

Richardson CD, Ray GJ, DeWitt MA, Curie GL, Corn JE. Enhancing homology-directed genome editing by catalytically active and inactive CRISPR-Cas9 using asymmetric donor DNA. Nat Biotechnol. 34(3) 339–44 (2016).

Shen MW, Arbab M, Hsu JY, Worstell D, Culbertson SJ, et al. Predictable and precise template-free CRISPR editing of pathogenic variants. Nature 563(7733) 646–651 (2018).

Shou J, Li J, Liu Y, Wu Q. Precise and Predictable CRISPR Chromosomal Rearrangements Reveal Principles of Cas9-Mediated Nucleotide Insertion. Mol. Cell 71(4) 498–509.e4 (2018).

Sternberg SH, Redding S, Jinek M, Greene EC, Doudna JA. DNA interrogation by the CRISPR RNA-guided endonuclease Cas9. Nature 507(7490) 62–67 (2014).

Sternberg SH, LaFrance B, Kaplan M, Doudna JA. Conformational control of DNA target cleavage by CRISPR–Cas9. Nature 527: 110–113 (2015).

Strick TR, Allemand JF, Bensimon D, Bensimon A, Croquette V. The elasticity of a single supercoiled DNA molecule. Science 271(5257): 1835–1837 (1996).

Strick TR, Croquette V, Bensimon D. Homologous pairing in stretched supercoiled DNA. Proc. Natl. Acad. Sci. (USA) 95(18): 10579–10583 (1998).

Szczelkun MD, Tikhomirova MS, Sinkunas T, Gasiunas G, Karvelis T, et al. Direct observation of R-loop formation by single RNA-guided Cas9 and Cascade effector complexes. PNAS 111(27) 9798–9803 (2014).

van den Broek B, Noom MC, Wuite GJ. DNA-tension dependence of restriction enzyme activity reveals mechanochemical properties of the reaction pathway. Nucleic Acids Res. 33(8) 2676–2684 (2005).

Wang Y, Mallon J, Wang H, Singh D, Hyun Jo M, Hua B, Bailey S, Ha T. Real-time observation of Cas9 postcatalytic domain motions. PNAS, 118(2) e2010650118 (2021).

Yang M, Peng S, Sun R, Lin J, Wang N, Chen C. The Conformational Dynamics of Cas9 Governing DNA Cleavage Are Revealed by Single-Molecule FRET. Cell Rep. 22(2) 372–382 (2018).

Zuo Z, Liu J. Cas9-catalyzed DNA Cleavage Generates Staggered Ends: Evidence from Molecular Dynamics Simulations. Sci. Rep. 5 37584 (2016).

