## Supplementary for "Shifted PAMs generate DNA overhangs and enhance SpCas9 post-catalytic complex dissociation"

**Extended Table 1**. "In-Del" and "5'-locating" data obtained by Sanger sequencing. The first column specifies the guide RNA used. The second column contains the sequencing results obtained after SpCas9-RNP cleavage, blunting and repair by T4 ligase; here the NGG PAM is shown in red, the nucleotide insertion (as specified) and deletion (specified as "-") are shown in green, and the number of times each event is observed, n, is specified. The third column contains the sequencing results obtained after SpCas9-RNP cleavage, Sbf-I cleavage, blunting and repair by T4 ligase (the Sbf-I end treated by Sbf-I and blunting kit to generate a double cytosine reference is shown in cyan, and "/" indicates the ligation site of the two broken ends).

| Guide RNAs | In-Del | 5'-locating |
|---|---|---|
| S2_T0 | TATTTGGGAATTCGGTACCTGG; n=53<br>TATTTGGGAATTCGGTTACCTGG; n=6<br>TATTTGGGAATTCGGT-CCTGG; n=1 | TGCATGCC/ACCTGG; n=29<br>TGCATGCC/TACCTGG; n=2 |
| S1_T0 | CCTTTTTATTTGGGAATTCGGT; n=37<br>CCTTTTTTATTTGGGAATTCGGT; n=1<br>CCTTTT-ATTTGGGAATTCGGT; n=1 | TTGCATGCC/TATTTGGGAATTCGGTACC; n=30<br>TTGCATGCC/TTTGGGAATTCGGTACC; n=1 |
| S1_T1 | CCTTTTTATTTGGGAATTCGGT; n=1<br>CCTTTT-ATTTGGGAATTCGGT; n=39 | TTGCATGCC/ATTTGGGAATTCGGTACC; n=34 |
| S1_T2 | CCTTTTTATTTGGGAATTCGGT; n=0<br>CCTTTT-ATTTGGGAATTCGGT; n=1<br>CCTTTT--TTTGGGAATTCGGT; n=52<br>CCTTTT----TGGGAATTCGGT; n=1 | TTGCATGCC/TTTGGGAATTCGGTACC; n=34 |
| S1_T3 | CCTTTTTATTTGGGAATTCGGT; n=16<br>CCTTTTTATTTTGGGAATTCGGT; n=5<br>CCTTTT-ATTTGGGAATTCGGT; n=2<br>CCTTTT--TTTGGGAATTCGGT; n=4<br>CCTTTT---TTGGGAATTCGGT; n=8 | TTGCATGCC/TTGGGAATTCGGTACC; n=42<br>TTGCATGCC/TGGGAATTCGGTACC; n=2<br>TTGCATGCC/ATTCGGTACC; n=1 |
| S1_T4 | CCTTTTTATTTGGGAATTCGGT; n=30<br>CCTTTTTATTTGTGGGAATTCGGT; n=1<br>CCTTTTTATTTTGGGAATTCGGT; n=5<br>CCTTTTTAT-TGGGAATTCGGT; n=1 | TTGCATGCC/TGGGAATTCGGTACC; n=35<br>TTGCATGCC/ATTCGGTACC; n=1 |
| S1_T-1 | CCTTTTTATTTGGGAATTCGGT; n=35<br>CCTTTT-ATTTGGGAATTCGGT; n=3<br>CCTTTT--TTTGGGAATTCGGT; n=1 | TTGCATGCC/TTATTTGGGAATTCGGTACC; n=35<br>TTGCATGCC/TATTTGGGAATTCGGTACC; n=2<br>TTGCATGCC/ATTTGGGAATTCGGTACC; n=3<br>TTGCATGCC/TTTGGGAATTCGGTACC; n=1 |
| S4_T2 | AGATTCATAAATTTGAGAGAGG; n=0<br>AGATTCATAAATTT-AGAGAGG; n=24<br>AGATTCATAAATTT--GAGAGG; n=13 | N.A. |
| S5_T2 | ATATCGAGGAGTTTAAATATGG; n=0<br>ATATCGAGGAGTTT-AATATGG; n=1<br>ATATCGAGGAGTTT--ATATGG; n=38 | TTGCATGCC/AATATGG; n=6<br>TTGCATGCC/ATATGG; n=31 |

**Extended Table 2.** Summary of mean lifetimes (or duration, DUR) obtained in single-molecule experiments.

| **$\langle t_{R\text{-}on} \rangle$** | | | | |
|---|---|---|---|---|
| **Conditions** | **Supercoil** | **Counts #** | **DUR (s)** | **Deviation (s)** |
| 1nM S1_T0 dSpCas9 | N8 | 138 | 9.6 | 1.0 |
| 1nM S1_T1 dSpCas9 | N8 | 141 | 10.7 | 1.1 |
| 1nM S1_T2 dSpCas9 | N8 | 300 | 56.1 | 4.1 |
| 1nM S2_T0 dSpCas9 | N8 | 189 | 40.3 | 3.5 |
| 1nM S1_T0 SpCas9 | N8 | 88 | 16.3 | 2.5 |
| 1nM S1_T1 SpCas9 | N8 | 81 | 27.9 | 4.0 |
| 1nM S1_T2 SpCas9 | N8 | 62 | 155 | 33.4 |
| 1nM S2_T0 SpCas9 | N8 | 72 | 83.0 | 14.8 |
| 1nM S2_T0 SpCas9 +RNAP | N8 | 67 | 89.0 | 16.3 |
| 1nM S3_T0 SpCas9 | N8 | 56 | 49.0 | 8.7 |
| 1nM S3_T0 SpCas9 +RNAP | N8 | 98 | 69.0 | 9.5 |
| 1nM S5_T0 SpCas9 | N8 | 49 | 46 | 10 |
| 1nM S5_T1 SpCas9 | N8 | 52 | 152 | 32 |
| 1nM S5_T2 SpCas9 | N8 | 38 | 1800 | 280 |
| 1nM S1_T0 SpCas9(D10A) | N8 | 58 | 10.8 | 2.0 |
| 1nM S1_T1 SpCas9(D10A) | N8 | 60 | 10.3 | 1.9 |
| 1nM S1_T2 SpCas9(D10A) | N8 | 78 | 66.4 | 10.8 |
| 1nM S2_T0 SpCas9(D10A) | N8 | 69 | 34.4 | 5.4 |
| 1nM S1_T0 SpCas9(H840A) | N8 | 82 | 206 | 32 |
| 1nM S1_T1 SpCas9(H840A) | N8 | 55 | 995 | 219 |
| 10nM S1_T2 SpCas9(H840A) | N8 | 58 | 1770 | 220 |
| 1nM S2_T0 SpCas9(H840A) | N8 | 65 | 929 | 149 |
| **$\langle t_{R\text{-}off} \rangle$** | | | | |
| **Conditions** | **Supercoil** | **Counts #** | **DUR (s)** | **Deviation (s)** |
| 1nM S1_T0 dSpCas9 | P5 | 68 | 17.6 | 4.4 |
| 1nM S1_T1 dSpCas9 | P5 | 160 | 86.6 | 10.8 |
| 1nM S1_T2 dSpCas9 | P5 | 137 | 64.6 | 10.7 |
| 1nM S2_T0 dSpCas9 | P5 | 75 | 23.0 | 5.1 |
| **$\langle t_{Super\text{-}off} \rangle$** | | | | |
| **Conditions** | **Counts #** | **DUR (s)** | **Deviation (s)** | **P*>3600s (%)** |
| 1nM S1_T0 SpCas9 | 93 | 29.7 | 4.0 | 2.2 |
| 1nM S1_T1 SpCas9 | 87 | 9.4 | 1.4 | 0 |
| 1nM S1_T2 SpCas9 | 69 | 5.5 | 1.2 | 10.1 |
| 1nM S2_T0 SpCas9 | 76 | 6.5 | 1.1 | 3.9 |
| 1nM S2_T0 SpCas9 +RNAP | 70 | 9.5 | 1.9 | 2.9 |
| 1nM S3_T0 SpCas9 | 59 | 35.2 | 6.8 | 0 |

| | | | | |
|---|---|---|---|---|
| 1nM S3_T0 SpCas9 +RNAP | 103 | 26.3 | 3.6 | 0 |
| 1nM S5_T0 SpCas9 | 47 | 3.2 | 0.8 | 0 |
| 1nM S5_T1 SpCas9 | 52 | 4.6 | 1.2 | 0 |
| 1nM S5_T2 SpCas9 | 37 | 2.7 | 1.6 | 48.6 |
| 1nM S1_T1 SpCas9(D10A) | 68 | 106 | 21 | 0 |
| 1nM S1_T2 SpCas9(D10A) | 82 | 59.0 | 13.2 | 28 |
| 1nM S1_T0 SpCas9(D10A) | 59 | 10230# | 940# | 81.4 |
| 1nM S2_T0 SpCas9(D10A) | 63 | 6750# | 370# | 84.1 |
| 1nM S1_T0 SpCas9(H840A) | 81 | 5.2 | 1.4 | 34.6 |
| 1nM S1_T1 SpCas9(H840A) | 55 | 2.5 | 0.6 | 16.4 |
| 10nM S1_T2 SpCas9(H840A) | 57 | 1.4 | 0.5 | 24.6 |
| 1nM S2_T0 SpCas9(H840A) | 65 | 4.1 | 1.0 | 26.2 |

P* > 3600s, the percentage of events last more than 3600 seconds

### Raw average elapse of supercoil maintained after R-loop formation given measured time-window and corresponding Deviation (SEM), due to majority of population last > 3600 seconds.

**$\langle t_{\text{Bead-off}} \rangle$**

| **Conditions** | **Counts #** | **DUR (s)** | **Deviation (s)** | **P*>7200s (%)** |
|---|---|---|---|---|
| 1nM S1_T0 SpCas9 | 93 | 9270# | 750# | 53.8 |
| 1nM S1_T1 SpCas9 | 87 | 490 | 104 | 6.9 |
| 1nM S1_T2 SpCas9 | 63 | 105 | 21 | 6.3 |
| 1nM S2_T0 SpCas9 | 76 | 16390# | 1140# | 73.7 |
| 1nM S2_T0 SpCas9 +RNAP | 69 | 884 | 271 | 43.5 |
| 1nM S3_T0 SpCas9 | 59 | 13690# | 920# | 78 |
| 1nM S3_T0 SpCas9 +RNAP | 103 | 282 | 69 | 47.6 |
| 1nM S5_T0 SpCas9 | 42 | 10090# | 610# | 83.3 |
| 1nM S5_T1 SpCas9 | 52 | 153 | 34 | 11.5 |
| 1nM S5_T2 SpCas9 | 19 | 33 | 15 | 15.8 |

P* > 7200s, the percentage of events last more than 7200 seconds.

### Raw average elapse of the incised molecules maintained given measured time-window and corresponding Deviation (SEM), due to majority of population last > 7200 seconds.

**Extended Table 3**. Guide RNA oligos used for ensemble biochemistry and single-molecule assays.

| **RNA_Name** | **Sequence (5'-3')** |
|---|---|
| tracrRNA | GGAACCAUUCAAAACAGCAUAGCAAGUUAAAAUAAGGCUAGUCCGUUAUCAACUUGA AAAAGUGGCACCGAGUCGGUGCUUUUUUU |
| crRNA S1_T0 | UACCGAAUUCCCAAAUAAAAGUUUUAGAGCUAUGCUGUUUUG |
| crRNA S1_T1 | GUACCGAAUUCCCAAAUAAAGUUUUAGAGCUAUGCUGUUUUG |
| crRNA S1_T2 | GGUACCGAAUUCCCAAAUAAGUUUUAGAGCUAUGCUGUUUUG |
| crRNA S1_T3 | AGGUACCGAAUUCCCAAAUAGUUUUAGAGCUAUGCUGUUUUG |
| crRNA S1_T4 | CAGGUACCGAAUUCCCAAAUGUUUUAGAGCUAUGCUGUUUUG |
| crRNA S1_T-1 | ACCGAAUUCCCAAAUAAAAAGUUUUAGAGCUAUGCUGUUUUG |
| crRNA S1_T-2 | CCGAAUUCCCAAAUAAAAAGGUUUUAGAGCUAUGCUGUUUUG |
| crRNA S2_T0 | UUAUUUGGGAAUUCGGUACCGUUUUAGAGCUAUGCUGUUUUG |
| crRNA S3_T0 | UAAUCGCAGGCCUUUUUAUUGUUUUAGAGCUAUGCUGUUUUG |
| crRNA S4_T2 | AAUAGAUUCAUAAAUUUGAGGUUUUAGAGCUAUGCUGUUUUG |
| crRNA S5_T0 | GAUAUCGAGGAGUUUAAAUAGUUUUAGAGCUAUGCUGUUUUG |
| crRNA S5_T1 | CGAUAUCGAGGAGUUUAAAUGUUUUAGAGCUAUGCUGUUUUG |
| crRNA S5_T2 | UCGAUAUCGAGGAGUUUAAAGUUUUAGAGCUAUGCUGUUUUG |
| crRNA S6_T2 | UCGGUACCUGGAGGAAGCCGGUUUUAGAGCUAUGCUGUUUUG |
| Seq_Kpn-80 | ACTACGGTTTCACCCTCTCCACCACC |
| Seq_Kpn+70 | CCATCTCGTAGGCCTGCTCAATC |

**Extended Table 4**. DNA oligos used for point mutagenesis and Sanger sequencing.

| Oligo Name | Sequence (5'-3') | Function |
|---|---|---|
| S1_PAM_mut_P1 | GTAATCGCAGGAGTTTTTATTTGGGAATTCGGTACC | Mutagenesis |
| S1_PAM_mut_P2 | CAAATAAAAACTCCTGCGATTACCAGCAGGCCTG | Mutagenesis |
| S4_PAM_mut1_P1 | TAAATTTGAGACAGGAGTCCCTAGTATGCATCG | Mutagenesis |
| S4_PAM_mut1_P2 | CTAGGGACTCCTGTCTCAAATTTATGAATCTATTATACAG | Mutagenesis |
| S4_PAM_mut2_P1 | ATTTGAGAGACTAGTCCCTAGTATGCATCGAATAG | Mutagenesis |
| S4_PAM_mut2_P2 | TACTAGGGACTAGTCTCTCAAATTTATGAATCTATTATACAG | Mutagenesis |
| S4_PAM_mut3_P1 | ATTTGAGACACTAGTCCCTAGTATGCATCGAATAG | Mutagenesis |
| S4_PAM_mut3_P2 | TACTAGGGACTAGTGTCTCAAATTTATGAATCTATTATACAG | Mutagenesis |
| S5_PAM_mut_P1 | GTTTAAATATCGCTGATGCATGAATTCGTTAATAACAGG | Mutagenesis |
| S5_PAM_mut_P2 | TCATGCATCAGCGATATTTAAACTCCTCGATATCGATTGG | Mutagenesis |
| S1_1st_A2T_P1 | GTAATCGCAGGCCTATTTATTTGGGAATTCGGTACC | Mutagenesis |
| S1_1st_A2T_P2 | CAAATAAATAGGCCTGCGATTACCAGCAGGCCTG | Mutagenesis |
| S1_1st_A2G_P1 | GTAATCGCAGGCCTCTTTATTTGGGAATTCGGTACC | Mutagenesis |
| S1_1st_A2G_P2 | CAAATAAAGAGGCCTGCGATTACCAGCAGGCCTG | Mutagenesis |
| S1_1st_A2C_P1 | GTAATCGCAGGCCTGTTTATTTGGGAATTCGGTACC | Mutagenesis |
| S1_1st_A2C_P2 | CAAATAAACAGGCCTGCGATTACCAGCAGGCCTG | Mutagenesis |
| S1_2nd_A2T_P1 | GTAATCGCAGGCCTTATTATTTGGGAATTCGGTACC | Mutagenesis |
| S1_2nd _A2T_P2 | CAAATAATAAGGCCTGCGATTACCAGCAGGCCTG | Mutagenesis |
| S1_2nd _A2G_P1 | GTAATCGCAGGCCTTCTTATTTGGGAATTCGGTACC | Mutagenesis |
| S1_2nd _A2G_P2 | CAAATAAGAAGGCCTGCGATTACCAGCAGGCCTG | Mutagenesis |
| S1_2nd _A2C_P1 | GTAATCGCAGGCCTTGTTATTTGGGAATTCGGTACC | Mutagenesis |
| S1_2nd _A2C_P2 | CAAATAACAAGGCCTGCGATTACCAGCAGGCCTG | Mutagenesis |
| Seq_Kpn-80 | ACTACGGTTTCACCCTCTCCACCACC | Sanger sequencing |
| Seq_Kpn+70 | CCATCTCGTAGGCCTGCTCAATC | Sanger sequencing |

**Extended Table 5**. Oligos used for cloning sgRNA genes into pX330-SpCas9 and pMCB320 vectors.

| **Oligo Name** | **Sequence (5'-3')** | **Function** |
|---|---|---|
| RAG1B_T0-O1 | CACCGACTTGTTTTCATTGTTCTC | Prey |
| RAG1B_T0-O2 | AAACGAGAACAATGAAAACAAGTC | Prey |
| RAG1B_T1-O1 | CACCGTGACTTGTTTTCATTGTTCT | Prey |
| RAG1B_T1-O2 | AAACAGAACAATGAAAACAAGTCAC | Prey |
| RAG1B_T2-O1 | CACCGATGACTTGTTTTCATTGTTC | Prey |
| RAG1B_T2-O2 | AAACGAACAATGAAAACAAGTCATC | Prey |
| RAG1C_T0-O1 | CACCGCACCTAACATGATATATTA | Prey |
| RAG1C_T0-O2 | AAACTAATATATCATGTTAGGTGC | Prey |
| RAG1C_T1-O1 | CACCGAGCACCTAACATGATATATT | Prey |
| RAG1C_T1-O2 | AAACAATATATCATGTTAGGTGCTC | Prey |
| RAG1C_T2-O1 | CACCGCAGCACCTAACATGATATAT | Prey |
| RAG1C_T2-O2 | AAACATATATCATGTTAGGTGCTGC | Prey |
| **Bait-D-O1/** RAG1D_T0-O1 | CACCGACCTTAAGGTTTTTGTGGA | **Bait** / Prey |
| **Bait-D-O2/** RAG1D_T0-O2 | AAACTCCACAAAAACCTTAAGGTC | **Bait** / Prey |
| RAG1D_T1-O1 | CACCGAGACCTTAAGGTTTTTGTGG | Prey |
| RAG1D_T1-O2 | AAACCCACAAAAACCTTAAGGTCTC | Prey |
| RAG1D_T2-O1 | CACCGAAGACCTTAAGGTTTTTGTG | Prey |
| RAG1D_T2-O2 | AAACCACAAAAACCTTAAGGTCTTC | Prey |
| **Bait-A-O1/** RAG1A_T0-O1 | CACCGCCTCTTTCCCACCCACCTT | **Bait** |
| **Bait-A-O2/** RAG1A_T0-O2 | AAACAAGGTGGGTGGGAAAGAGGC | **Bait** |
| RAG1K_T0-O1 | CACCGTAGTACCAAGCTCCTTGCAG | Prey |
| RAG1K_T0-O2 | AAACCTGCAAGGAGCTTGGTACTAC | Prey |
| RAG1K_T1-O1 | CACCGTTAGTACCAAGCTCCTTGCA | Prey |
| RAG1K_T1-O2 | AAACTGCAAGGAGCTTGGTACTAAC | Prey |
| RAG1K_T2-O1 | CACCGCTTAGTACCAAGCTCCTTGC | Prey |
| RAG1K_T2-O2 | AAACGCAAGGAGCTTGGTACTAAGC | Prey |
| **Bait-L-O1/** RAG1L_T0-O1 | CACCGACACATTCTGGCTGACCCTG | **Bait** / Prey |
| **Bait-L-O2/** RAG1L_T0-O2 | AAACCAGGGTCAGCCAGAATGTGTC | **Bait** / Prey |
| RAG1L_T1-O1 | CACCGAACACATTCTGGCTGACCCT | Prey |
| RAG1L_T1-O2 | AAACAGGGTCAGCCAGAATGTGTTC | Prey |
| RAG1L_T2-O1 | CACCGGAACACATTCTGGCTGACCC | Prey |

| | | |
|---|---|---|
| RAG1L_T2-O2 | AAACGGGTCAGCCAGAATGTGTTCC | Prey |
| RAG1M_T0-O1 | CACCGTCCATGCTTCCCTACTGACC | Prey |
| RAG1M_T0-O2 | AAACGGTCAGTAGGGAAGCATGGAC | Prey |
| RAG1M_T1-O1 | CACCGATCCATGCTTCCCTACTGAC | Prey |
| RAG1M_T1-O2 | AAACGTCAGTAGGGAAGCATGGATC | Prey |
| RAG1M_T2-O1 | CACCGTATCCATGCTTCCCTACTGA | Prey |
| RAG1M_T2-O2 | AAACTCAGTAGGGAAGCATGGATAC | Prey |
| RAG1N_T0-O1 | CACCGGTGCAATGAGGAGGTCAGTT | Prey |
| RAG1N_T0-O2 | AAACAACTGACCTCCTCATTGCACC | Prey |
| RAG1N_T1-O1 | CACCGAGTGCAATGAGGAGGTCAGT | Prey |
| RAG1N_T1-O2 | AAACACTGACCTCCTCATTGCACTC | Prey |
| RAG1N_T2-O1 | CACCGGAGTGCAATGAGGAGGTCAG | Prey |
| RAG1N_T2-O2 | AAACCTGACCTCCTCATTGCACTCC | Prey |
| RAG1O_T0-O1 | CACCGCTCGGAGAGCTCAGAAGCAC | Prey |
| RAG1O_T0-O2 | AAACGTGCTTCTGAGCTCTCCGAGC | Prey |
| RAG1O_T1-O1 | CACCGACTCGGAGAGCTCAGAAGCA | Prey |
| RAG1O_T1-O2 | AAACTGCTTCTGAGCTCTCCGAGTC | Prey |
| RAG1O_T2-O1 | CACCGGACTCGGAGAGCTCAGAAGC | Prey |
| RAG1O_T2-O2 | AAACGCTTCTGAGCTCTCCGAGTCC | Prey |
| GFP-CTR-O1 | TTGGACACATTCTGGCTGACCCTGGTTTAAGAGC | Control |
| GFP-CTR-O2 | TTAGCTCTTAAACCAGGGTCAGCCAGAATGTGTCCAACAAG | Control |
| GFP_T0-O1 | TTGGCTGCACGCCGTAGGTCAGGGGTTTAAGAGC | GFP-KO |
| GFP_T0-O2 | TTAGCTCTTAAACCCCTGACCTACGGCGTGCAGCCAACAAG | GFP-KO |
| GFP_T1-O1 | TTGGACTGCACGCCGTAGGTCAGGGTTTAAGAGC | GFP-KO |
| GFP_T1-O2 | TTAGCTCTTAAACCCTGACCTACGGCGTGCAGTCCAACAAG | GFP-KO |
| GFP_T2-O1 | TTGGCACTGCACGCCGTAGGTCAGGTTTAAGAGC | GFP-KO |
| GFP_T2-O2 | TTAGCTCTTAAACCTGACCTACGGCGTGCAGTGCCAACAAG | GFP-KO |

**Extended Table 6**. Oligos used for Nova-seq library preparation.

| Oligo name | Sequence (5'-3') |
|---|---|
| Bait-A-Bio | /5bioG/AGGACTGCTGGAGATTGCTC |
| Bait-L-Bio | /5bioG/TCAGCGACAGAAGATGTTGGC |
| Bait-D-Bio | /5bioG/CCAGCTAGTACAATGAGGCTAAT |
| Adapter-upper | GCGACTATAGGGCACGCGTGGTNNNNNN/3AmMO/ |
| Adapter-lower | /5Phos/ACCACGCGTGCCCTATAGTCGC/3AmMO/ |
| AP2I7 | TCTCGGCATTCCTGCTGAACCGCTCTTCCGATCTGACTATAGGGCACGCGTGGT |
| AP2I7-novo | GTGACTGGAGTTCAGACGTGTGCTCTTCCGATCTGACTATAGGGCACGCGTGGT |
| P7I7 | CAAGCAGAAGACGGCATACGAGATCGGTCTCGGCATTCCTGCTGAACC |
| P7I7-i12 | CAAGCAGAAGACGGCATACGAGATAGTACAAGGTGACTGGAGTTCAGACGTG |
| P7I7-i36 | CAAGCAGAAGACGGCATACGAGATCCAGTTCAGTGACTGGAGTTCAGACGTG |
| P7I7-i53 | CAAGCAGAAGACGGCATACGAGATGCTCGGTAGTGACTGGAGTTCAGACGTG |
| P5I5 | AATGATACGGCGACCACCGAGATCTACACTCTTTCCCTACACGACGCT |
| P5I5-i11 | AATGATACGGCGACCACCGAGATCTACACAACGTGATACACTCTTTCCCTACACGACG |
| P5I5-i37 | AATGATACGGCGACCACCGAGATCTACACCCGAAGTAACACTCTTTCCCTACACGACG |
| P5I5-i54 | AATGATACGGCGACCACCGAGATCTACACGGAGAACAACACTCTTTCCCTACACGACG |
| I5-Bait-A-02 | ACTCTTTCCCTACACGACGCTCTTCCGATCTTGGAGAGGGTTTCCCCTCAAAG |
| I5-Bait-A-03 | ACTCTTTCCCTACACGACGCTCTTCCGATCTAAGGAGAGGGTTTCCCCTCAAAG |
| I5-Bait-A-04 | ACTCTTTCCCTACACGACGCTCTTCCGATCTGCTCGAGAGGGTTTCCCCTCAAAG |
| I5-Bait-A-05 | ACTCTTTCCCTACACGACGCTCTTCCGATCTGCCATGAGAGGGTTTCCCCTCAAAG |
| I5-Bait-L-01 | ACACTCTTTCCCTACACGACGCTCTTCCGATCTATTCAGCATGGATATCGGCAAGAGGG |
| I5-Bait-L-02 | ACACTCTTTCCCTACACGACGCTCTTCCGATCTACTTGGCATGGATATCGGCAAGAGGG |
| I5-Bait-L-03 | ACACTCTTTCCCTACACGACGCTCTTCCGATCTGTGTTGCATGGATATCGGCAAGAGGG |
| I5-Bait-L-04 | ACACTCTTTCCCTACACGACGCTCTTCCGATCTTCGAGGCATGGATATCGGCAAGAGGG |
| I5-Bait-D-02 | ACTCTTTCCCTACACGACGCTCTTCCGATCTAAGGAGCACCTAATGTATACTGGG |
| I5-Bait-D-03 | ACTCTTTCCCTACACGACGCTCTTCCGATCTGCTCGAGCACCTAATGTATACTGGG |
| I5-Bait-D-04 | ACTCTTTCCCTACACGACGCTCTTCCGATCTGCCATGAGCACCTAATGTATACTGGG |
| I5-Bait-D-05 | ACTCTTTCCCTACACGACGCTCTTCCGATCTCGAGAGAGCACCTAATGTATACTGGG |
| I5-Bait-D-06 | ACTCTTTCCCTACACGACGCTCTTCCGATCTGTGTTGAGCACCTAATGTATACTGGG |
| I5-Bait-D-07 | ACTCTTTCCCTACACGACGCTCTTCCGATCTACTTGGAGCACCTAATGTATACTGGG |
| I5-Bait-D-08 | ACTCTTTCCCTACACGACGCTCTTCCGATCTAGTGAGAGCACCTAATGTATACTGGG |
| I5-Bait-D-09 | ACTCTTTCCCTACACGACGCTCTTCCGATCTTCGAAGAGCACCTAATGTATACTGGG |

**Extended Table 7**. Information for libraries obtained by Nova-seq.

| Library | Bait Breaks | Replicate | Raw Reads | Normalized Reads | Total Junctions | Junctions in Prey Region | | | | |
|---|---|---|---|---|---|---|---|---|---|---|
| Fig.1e | | | | | | | | | | |
| | | | | | | RAG1B | RAG1C | RAG1D | | |
| L-Ctr | Cas9:Bait-L | #1 | 5511798 | 5511798 | 80004 | 11 | 10 | 12 | | |
| | | #2 | 6818869 | 5866397 | 157147 | 7 | 2 | 5 | | |
| L-T0s | Cas9:Bait-L | #1 | 6427963 | 5511798 | 25728 | 209 | 234 | 320 | | |
| | | #2 | 8873803 | 5866397 | 44705 | 211 | 192 | 283 | | |
| L-T1s | Cas9:Bait-L | #1 | 5882578 | 5511798 | 38730 | 8 | 125 | 185 | | |
| | | #2 | 7066535 | 5866397 | 74187 | 1 | 119 | 149 | | |
| L-T2s | Cas9:Bait-L | #1 | 7281314 | 5511798 | 52736 | 9 | 13 | 13 | | |
| | | #2 | 5866397 | 5866397 | 44287 | 8 | 2 | 4 | | |
| Fig.1f | | | | | | | | | | |
| | | | | | | RAG1B | RAG1C | RAG1D | | |
| A-Ctr | Cas9:Bait-A | #1 | 9434312 | 8671237 | 304209 | 171 | 81 | 72 | | |
| | | #2 | 11973200 | 9701292 | 280144 | 166 | 68 | 66 | | |
| A-T0s | Cas9:Bait-A | #1 | 10908226 | 8671237 | 10800 | 242 | 397 | 435 | | |
| | | #2 | 9701292 | 9701292 | 7109 | 207 | 350 | 376 | | |
| A-T1s | Cas9:Bait-A | #1 | 8928036 | 8671237 | 34384 | 63 | 326 | 471 | | |
| | | #2 | 11225474 | 9701292 | 49401 | 42 | 353 | 601 | | |
| A-T2s | Cas9:Bait-A | #1 | 8671237 | 8671237 | 78487 | 89 | 58 | 106 | | |
| | | #2 | 11100144 | 9701292 | 75456 | 64 | 37 | 63 | | |
| Fig S5 | | | | | | | | | | |
| | | | | | | RAG1K | RAG1L | RAG1M | RAG1N | RAG1O |
| D-Ctr | Cas9:Bait-D | #1 | 12020563 | 8816431 | 303462 | 6 | 7 | 4 | 3 | 3 |
| | | #2 | 8432895 | 8432895 | 341509 | 6 | 9 | 7 | 5 | 3 |
| D-T0s | Cas9:Bait-D | #1 | 10290299 | 8816431 | 49486 | 336 | 106 | 101 | 191 | 457 |
| | | #2 | 10891188 | 8432895 | 33081 | 273 | 94 | 73 | 129 | 360 |
| D-T1s | Cas9:Bait-D | #1 | 9521505 | 8816431 | 115671 | 150 | 321 | 375 | 424 | 332 |
| | | #2 | 10687608 | 8432895 | 121161 | 176 | 392 | 460 | 592 | 432 |
| D-T2s | Cas9:Bait-D | #1 | 8816431 | 8816431 | 111189 | 2 | 28 | 1 | 1 | 2 |
| | | #2 | 9963500 | 8432895 | 134497 | 1 | 36 | 6 | 4 | 9 |

Bed of Prey Region

| Prey | Chr | Start | End |
|---|---|---|---|
| RAG1B | chr11 | 36594810 | 36594840 |
| RAG1C | chr11 | 36594681 | 36594711 |
| RAG1D | chr11 | 36594560 | 36594590 |
| RAG1K | chr11 | 36595678 | 36595709 |
| RAG1L | chr11 | 36595748 | 36595779 |
| RAG1M | chr11 | 36595860 | 36595891 |
| RAG1N | chr11 | 36595943 | 36595974 |
| RAG1O | chr11 | 36596070 | 36596101 |

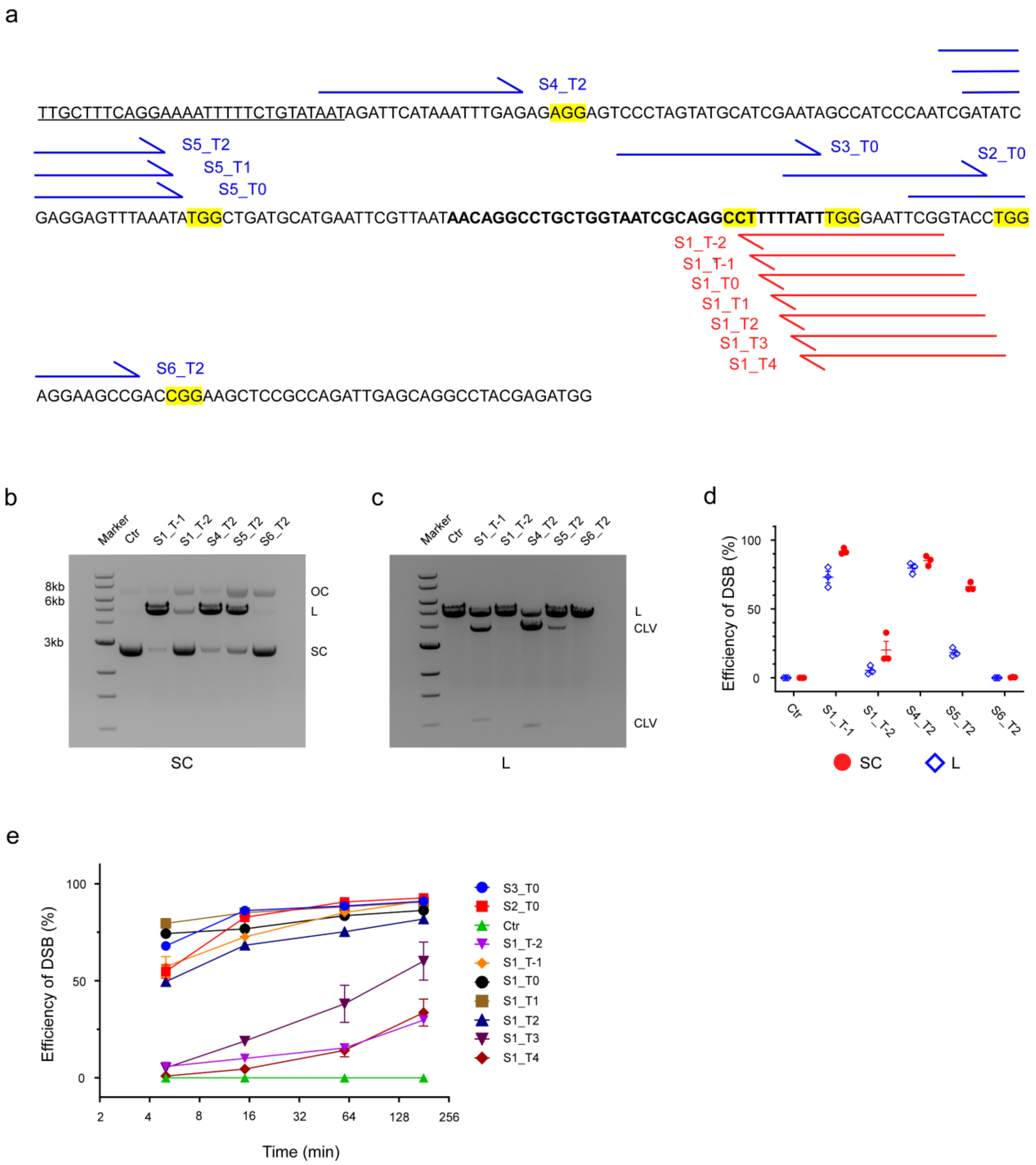

**Extended Data Fig.1 Cleaving DNA substrates using SpCas9-coupled guides that recognize canonical and shifted PAMs. (a)** Guide RNAs were designed to target a bacterial transcript (shown 5'-3') that contains an N25 promoter (underlined), a transcribed region and a tR2 terminator (bold). The 5'→3' blue and red arrows represent the 20 bp protospacers located on the template and non-template strands of the bacterial transcript, respectively. The corresponding PAMs are highlighted in yellow. **(b-d)** S1_T1, S1_T2, S4_T2, S5_T2, and S6_T2 SpCas9 RNPs that generate **(b)** open circular (OC) and linear (L) DNA on supercoiled (SC) DNA substrates, and **(c)** two cleaved (CLV) fragments on linearized DNA substrates. **(d)** Efficiencies of DSB formation on supercoiled circular (SC) and linear (L) DNA are quantified and normalized. **(e)** Ensemble assay of SpCas9 with various guides on supercoiled DNA substrates. Cleaved samples were taken at 5, 15, 60 and 180 min time points, loaded on an agarose gel and quantified by ImageJ. Means and corresponding SEMs are shown in the plot.

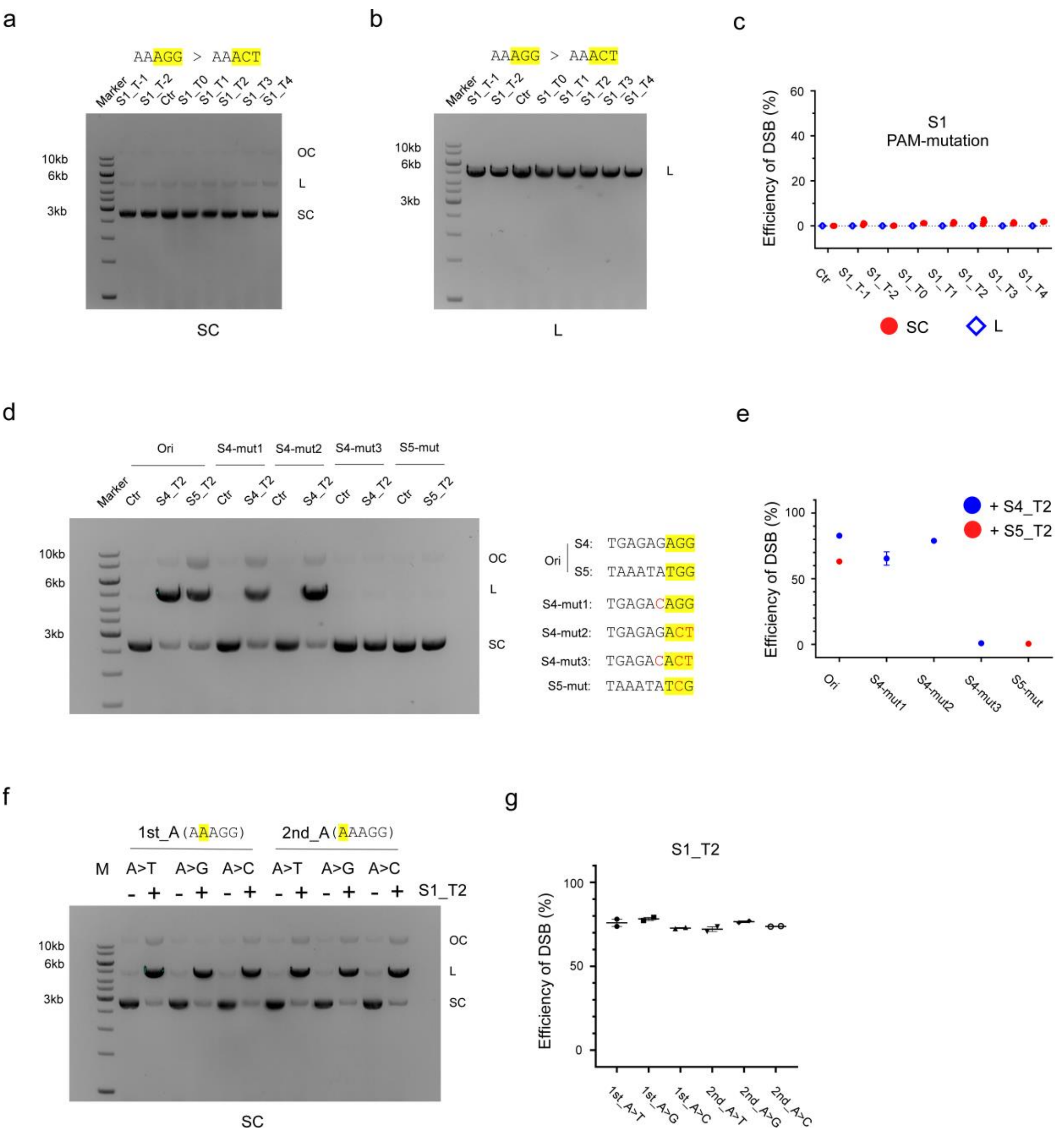

**Extended Data Fig.2 Different mutations were introduced into PAM or PAM-adjacent sites to assess the impact on SpCas9 cleavage activity using various guides. (a-c)** The AGG PAM of the S1 locus was mutated to ACC and supercoiled DNA ('SC', a) or linearized DNA ('L', b) bearing this mutation was cleaved by S1_T1, S1_T2, S1_T0, S1_T1, S1_T2, S1_T3 and S1_T4 SpCas9, respectively. Cleavage efficiencies were quantified, normalized, and plotted ©, with error bar (SEM). **(d-e)** Three mutants for the S4 locus were generated: S4-mut1 (PAM-adjacent guanine mutation), S4-mut2 (PAM mutation), and S4-mut3 (double mutation). One mutant for the S5 locus was generated, S5-mut. These mutants together with the original DNA (Ori) were cleaved by S4_T2 or S5_T2 SpCas9 (d), and the efficiency of DSB was calculated and normalized by subtracting the controls and shown in percentage with SEM (e). **(f-g)** The first and second adenines adjacent to the canonical PAM of the S1 locus were mutated to T, G, C respectively. The supercoiled forms of the corresponding mutants were cleaved by S1_T2 SpCas9 (f), and the efficiencies were calculated and shown in the right panel (g).

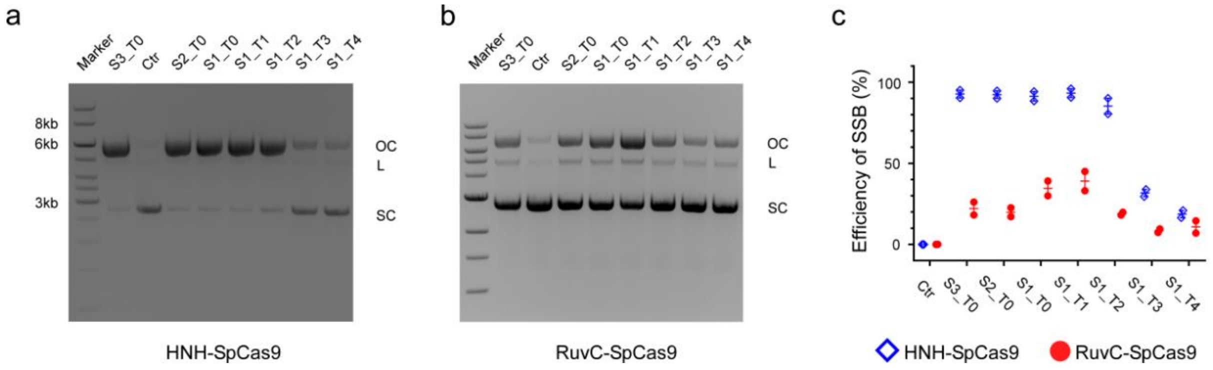

**Extended Data Fig.3 Supercoiled DNA with canonical and spaced PAMs were nicked using single-nuclease SpCas9 mutants. (a)** HNH-SpCas9 (D10A) nickase (containing an active HNH domain but inactive RuvC domain) was assembled with various dgRNAs into RNPs to incise the corresponding target double-stranded circular DNA containing either a canonical PAM or a shifted PAM. **(b)** As with the prior panel but using RuvC-SpCas9 (H840A) nickase (containing an active RuvC domain but inactive HNH domain). **(c)** Efficiencies of SSB formation are quantified, normalized and shown as blue diamonds for HNH-SpCas9(D10A) and red dots for RuvC-SpCas9(H840A). Error bars indicate SEM.

a

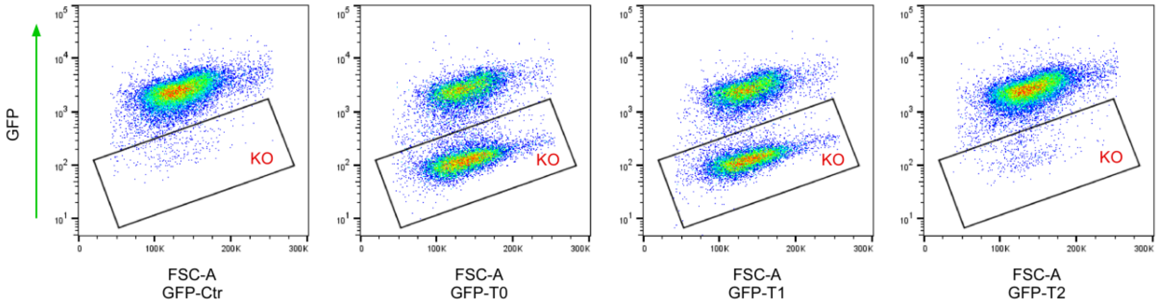

b

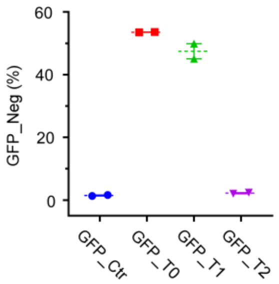

**Extended Data Fig.4 Canonical and PAM-shifted RNPs applied to gene knockout. (a)** Flow cytometry measurement of GFP in K562-iCas9-GFP cells nucleofected by pMCB320 bearing sgRNA genes of GFP-Ctr, GFP_T0, GFP_T1 and GFP_T2, respectively. **(b)** The percentages of GFP-negative cells, GFP_Neg ("KO" in **a**) are plotted.

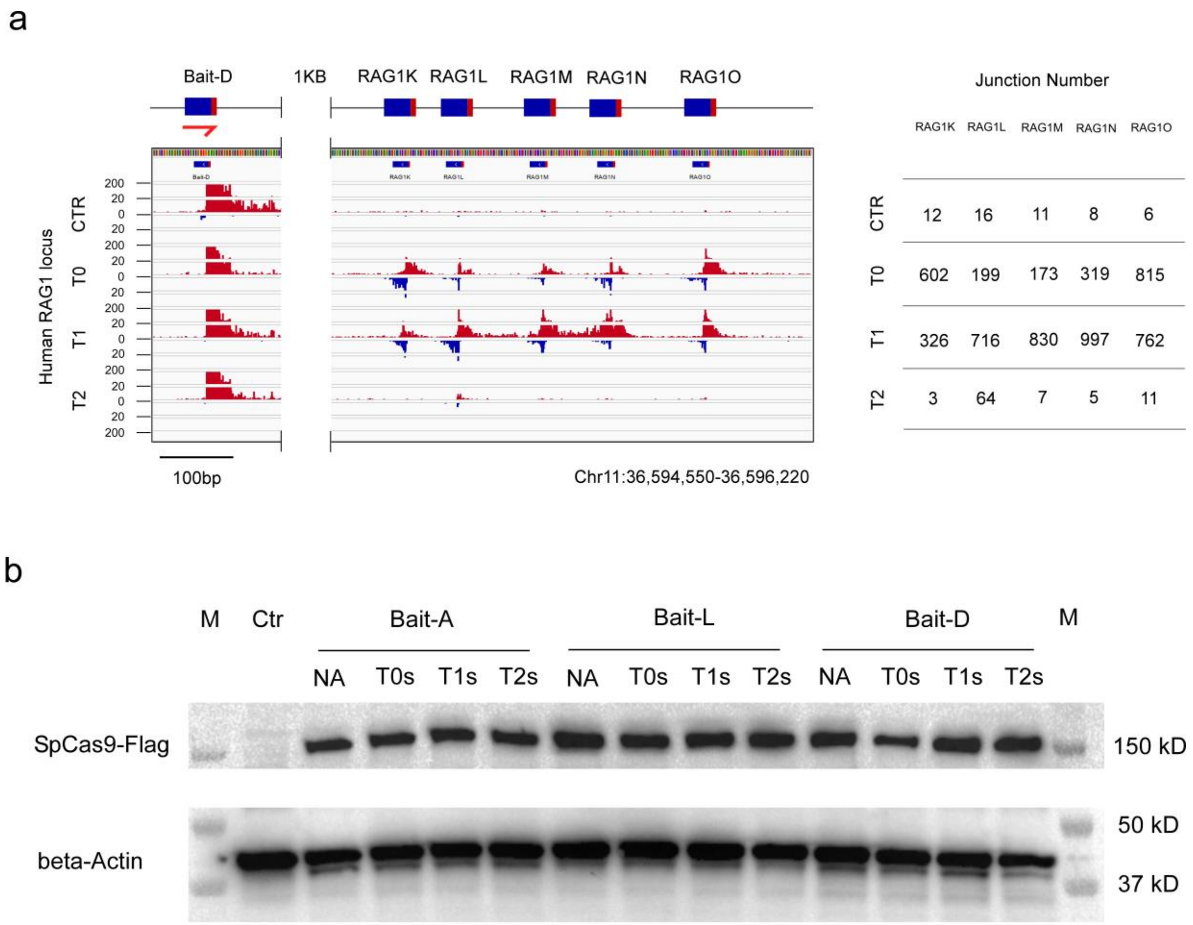

**Extended Data Fig.5 Comparing DSBs obtained using canonical and shifted-PAM guides at five additional sites.** (**a**) Bait-D (harpoon) targets a locus > 1kb upstream of the five prey sites RAG1K-O, and was delivered alone as negative control (CTR) or together with canonical PAM (T0) and shifted-PAM (T1, T2) sgRNAs into HEK293T cells, respectively. IGV-plot shows both T0 and T1 can generate decent amounts of cleavage at all five sites, while cleavage generated by T2 was barely detected excepted with the RAG1L_T2 guide. The number of junctions for each cluster is provided in the right panel. (**b**) Expression levels of flag-tagged SpCas9 (~150 kD) for all conditions including Bait-A, Bait-L and Bait-D series were determined by probing with an anti-flag antibody, and beta-actin (~40 kD) was used as a loading control.

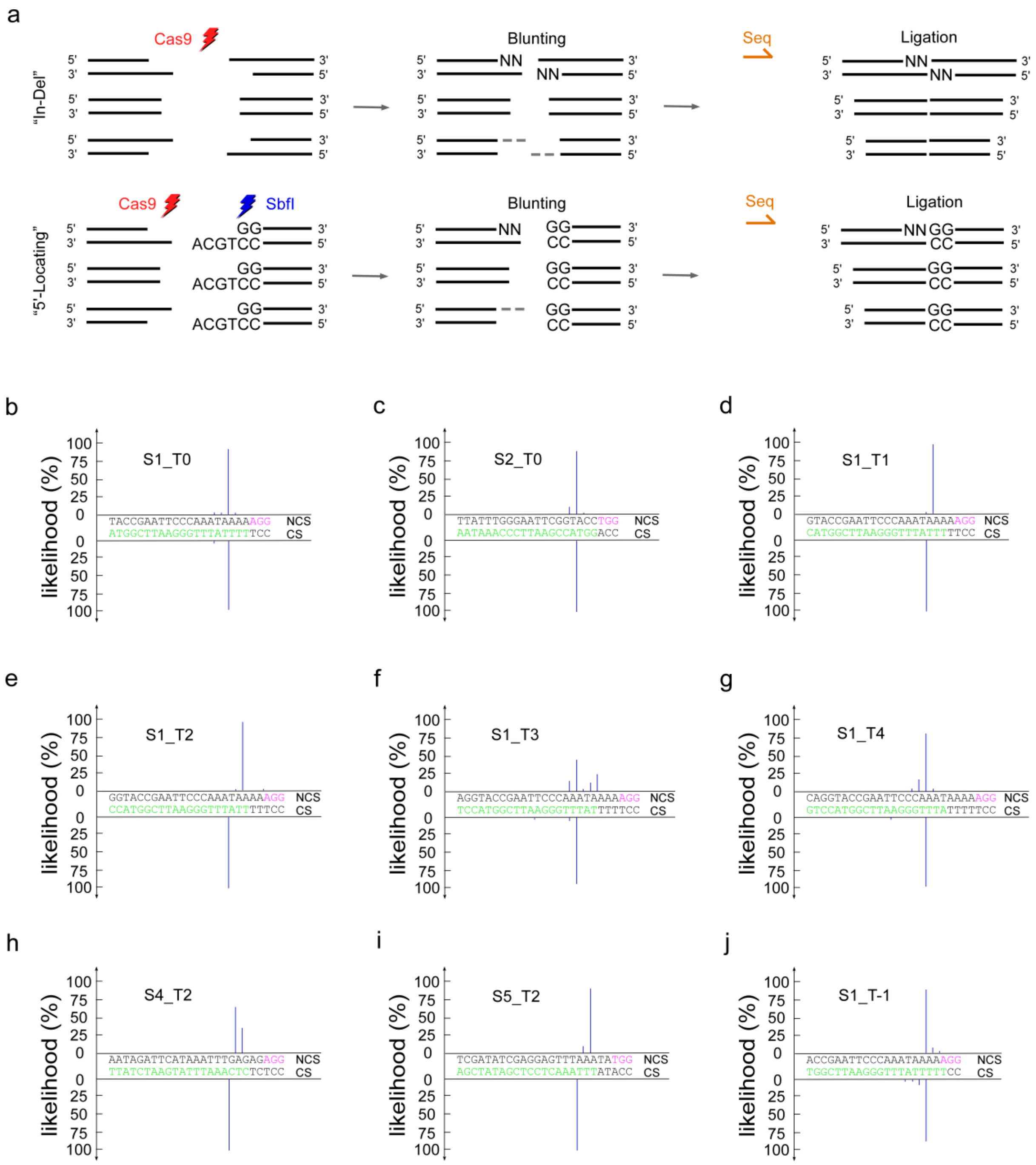

**Extended Data Fig.6 Identifying cleavage sites by "In-Del" and "5'-locating" approaches. (a)** Sketch of "In-Del" and "5'-locating" end-mapping methods, in which supercoiled DNA is processed by either Cas9 or by Cas9 followed by Sbf-I digestion and blunting (either by 5'→3' nucleotide addition at 3'-recessed ends or 3'→5' nucleotide deletion at 3'-overhang ends. DNA is then ligated and Sanger-sequenced (Seq; see Methods for details). **(b-j)** Resulting likelihood plots of cleavage sites for S1_T0, S2_T0, S1_T1, S1_T2, S1_T3, S1_T4, S4_T2, S5_T2 and S1_T-1 SpCas9. The height of the blue bar represents the likelihood of incision at the position of the DNA strand indicated. Green and magenta letters denote the protospacers and nearby canonical PAMs, respectively. At least 30 samples were sequenced for each condition (See Extend Table 1).

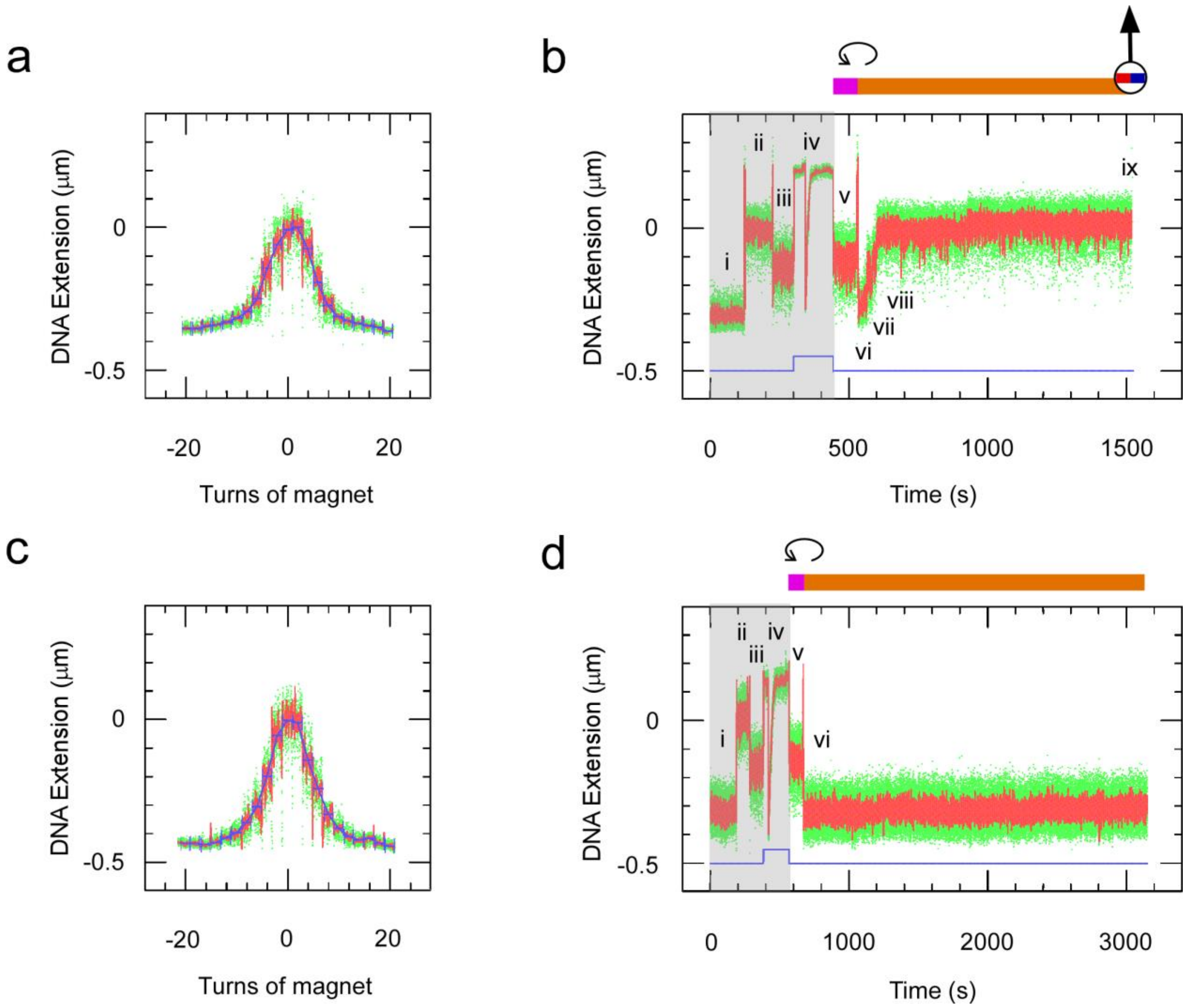

**Extended Data Fig.7 Procedure for DNA molecule calibration and dgRNA-SpCas9 injection using single-molecule magnetic trapping. (a)** Typical extension vs. supercoiling curve showing a DNA molecule rotated from 20 turns of positive supercoiling to 20 turns of negative supercoiling under a constant ~0.2 pN extending force, allowing us to identify un-nicked dsDNA molecules and also to calibrate the change in DNA extension for a unit change in DNA writhe (Strick et al. 1996), here typically ~55 nm/turn. **(b)** Extension time-trace for the molecule calibrated in (a), providing real-time references at a constant 0.2 pN extending force (blue line) for its extension at (i) 8 turns of negative supercoiling, (ii) zero supercoiling (ie the torsionally-relaxed state), and (iii) 5 turns of positive supercoiling before (iv) increasing the applied force to 1 pN and injecting S1_T0 SpCas9 RNPs (seen as a downward spike in extension as flow pushes the bead towards the surface). Once flow ends and the bead returns to its extension prior to injection (v) the applied force is returned to 0.2 pN and the extension is verified to return to that see in (iii). After a few minutes (vi) DNA molecules are returned to 8 turns of negative supercoiling. (vii) R-loop formation and (viii) supercoil loss are followed by (ix) terminal loss of the magnetic bead. **(c,d)** as in (a,b) but with injection of only the S1_T0 guide (ie without SpCas9) shows that no R-loops are formed within 2000 seconds of detection window. This experiment was repeated on a total of 72 molecules.

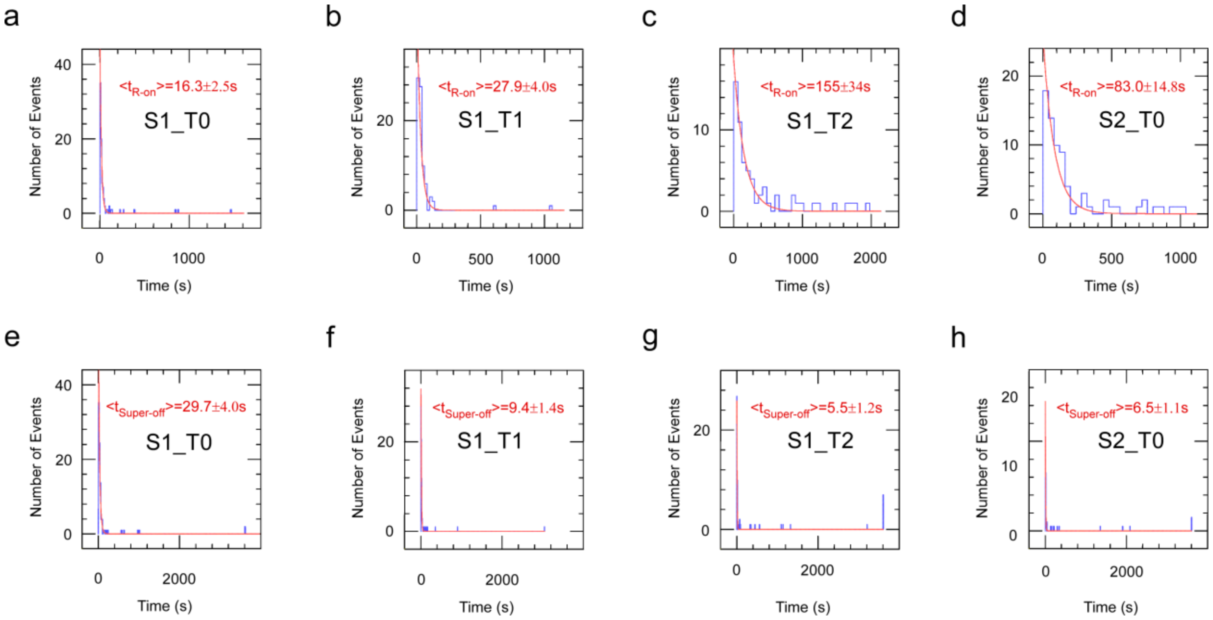

**Extended Data Fig.8 Characterization of the kinetics of R-loop formation and supercoiling loss using various dgRNA-SpCas9 RNPs.** (**a-d**) Distributions of waiting times for R-loop formation obtained for four different RNPs (blue) are fitted to single-exponential decays (red) to obtain the corresponding average waiting time, <$t_{R-on}$> (± SEM), for **(a)** S1_T0 SpCas9, **(b)** S1_T1 SpCas9, **(c)** S1_T2 SpCas9 and **(d)** S2_T0 SpCas9. (**e-h**) Similarly, the mean time elapsed between R-loop formation and supercoil dissociation, <$t_{Super-off}$>, was calculated for **(e)** S1_T0 SpCas9, **(f)** S1_T1 SpCas9, **(g)** S1_T2 SpCas9 and **(h)** S2_T0 SpCas9. See Extended Table 2 for further details.

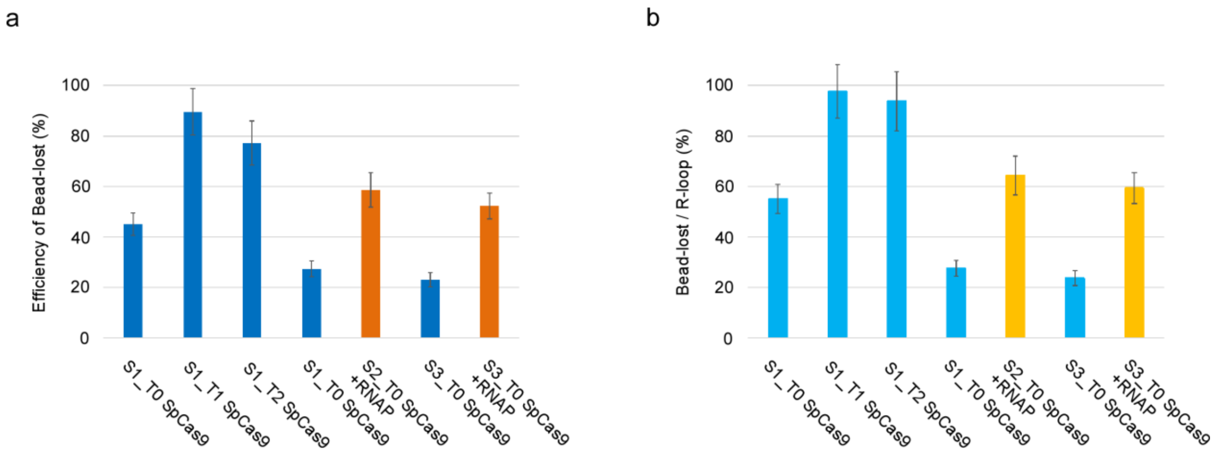

**Extended Data Fig.9 Apparent efficiency of DSB formation via magnetic trapping microscopy as determined by terminal loss of the magnetic bead within a three-hour window. (a)** Raw percentages of all observed DNA molecules displaying terminal bead loss in assays including S1_T0 SpCas9 (45.2 ± 6.6% SD or 47 of n=104 molecules), S1_T1 SpCas9 (89.5 ± 9.7% SD, or 85 of n=95 molecules), S1_T2 SpCas9 (77.2 ± 9.9% SD, or 61 of n=79 molecules), S2_T0 SpCas9 (27.3 ± 5.9% SD, 21 of n=77 molecules) and S3_T0 SpCas9 (23.1 ± 5.9% SD, or 15 of n=65 molecules) (blue bars). Experiments combining S2-T0 SpCas9 and RNAP (58.7 ± 8.8% SD, or 44 of n = 75 molecules), and S3_T0 SpCas9 and RNAP (52.3 ± 7.0% SD, or 56 of n=107 molecules) are also quantified (orange bars). **(b)** Normalized percentages of those R-loop-forming DNA molecules which go on to display terminal bead loss for: S1_T0 SpCas9 (55.1%, n=89), S1_T1 SpCas9 (97.7%, n=86), S1_T2 SpCas9 (93.8%, n=64), S2_T0 SpCas9 (27.6%, n=76) and S3_T0 SpCas9 (23.7%, n=59), are normalized by dividing the number of molecules that formed R-loops (cyan bars with SEMs). Similarly, the normalized percentages for S2_T0 SpCas9 plus RNAP (64.3%, n=70) and NT00-SpCas9 plus RNAP (59.4%, n=96) are shown in yellow bars.

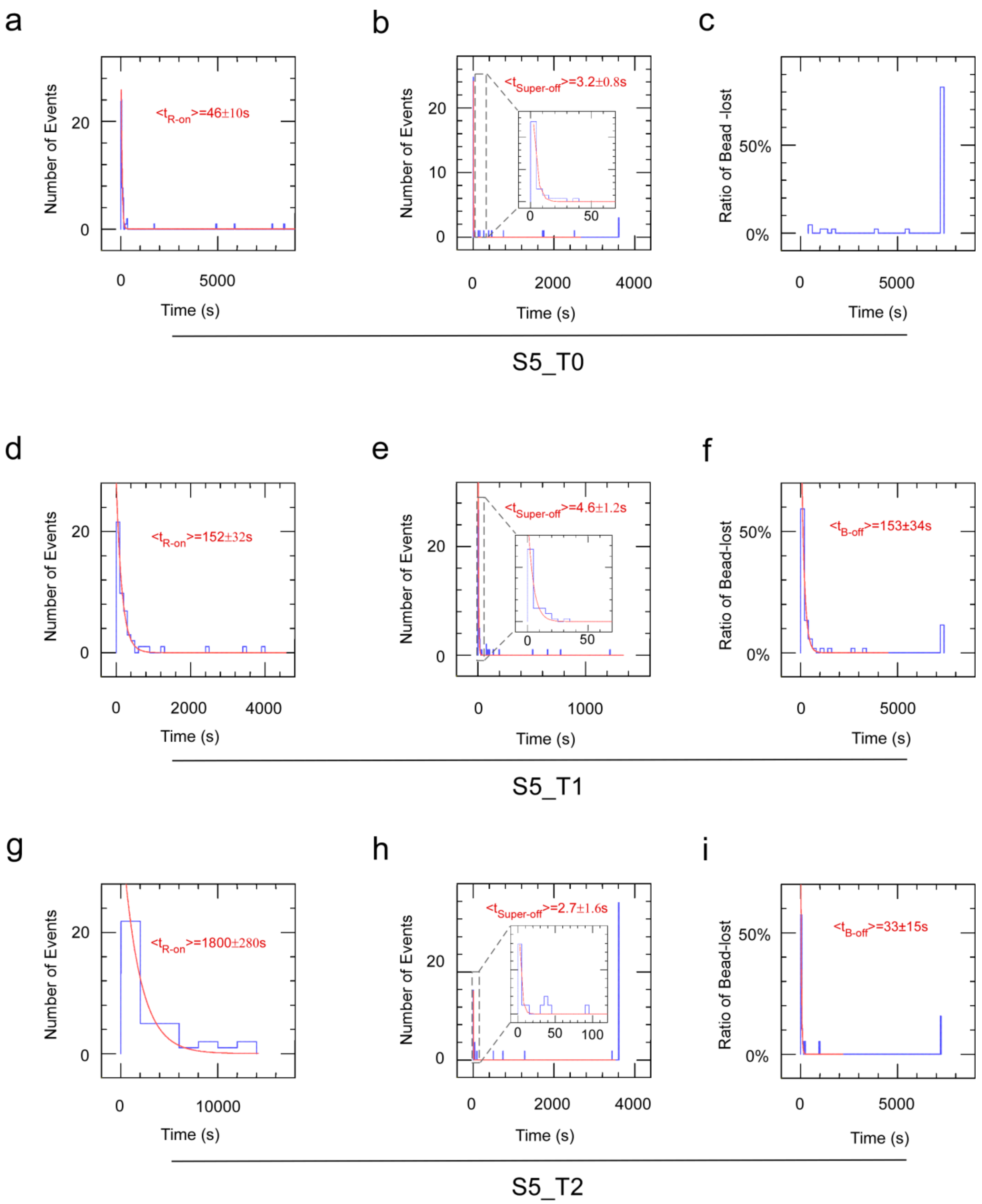

**Extended Data Fig.10 Comparing the kinetics of <$t_{R\text{-}on}$>, <$t_{Super\text{-}off}$> and <$t_{B\text{-}off}$> on extended site S5 using canonical and PAM-shifted guides.** (a-c) Canonical PAM S5_T0 RNPs generated R-loops within 46 ± 10s on average (n=49), caused loss of supercoiling within 3.2 ± 0.8s on average (n=47), and resulted in a long-lasting post-catalytic complex as 83.3% of molecules were broken within a 7200-second window. (d-f) Shifted PAM S5_T1 RNPs generated R-loops more slowly, i.e. within 152 ± 32s on average (n=52), caused loss of supercoiling within 4.6 ± 1.2s on average (n=52) and resulted in an unstable post-catalytic complex as 88.5% of molecules were broken within a 7200-second window. <$t_{B\text{-}off}$> was characterized as 153 ±34 s (n=52). (g-i) Shifted-PAM S5_T2 RNPs generated R-loops in the slowest manner, i.e. within 1800 ± 280s on average (n=38), caused loss of supercoiling within 2.7 ± 1.6s on average (n=37) and resulted in an unstable post-catalytic complex as 84.2% of molecules dissociated within an 7200 second window. $<t_{B-off}>$ was characterized as 33 ±15 s (n=19). See Extended Table 2 for specific parameters.
